# Spatial and network principles behind neural generation of locomotion

**DOI:** 10.1101/2024.10.03.616472

**Authors:** Salif Komi, August Winther, Grace A. Houser, Thomas Topilko, R.J.F. Sørensen, Silas Dalum Larsen, Madelaine C. Adamsson Bonfils, Guanghui Li, Rune W. Berg

## Abstract

Generation of locomotion is a fundamental function of the spinal cord, yet the underlying principles remain unclear. In particular, the relationship between neuronal cell types, networks and functions has been difficult to establish^1,2^. Here, we propose principles by which functions arise primarily from spatial features of the cord. First, we suggest that projections of distinct cell types constitute an asymmetrical “Mexican hat” topology, i.e. local excitation and surrounding inhibition with dissimilar length of projection along the rostro-caudal axis. Second, this projection topology constitutes the mechanism of rhythm- and pattern generation of mammalian locomotion. Third, the role of segregation of cell types in the transversal plane is for descending fibers to find appropriate targets. Modulation of these targets allows control of motor activity by adjusting the symmetry of the projection topology. We extract these principles via a model of the mouse spinal cord, where networks are constructed by probabilistic sampling of synaptic connections from cell-specific projection patterns, which are based on previous studies^3, 4^. The cell-type distributions are derived from single-cell RNA sequencing combined with spatial transcriptomics^5^. We find that essential aspects of locomotion are readily reproduced and controlled without requiring parameter optimization, and several experimental observations can now be explained mechanistically. Further, two main features are predicted: propagating bumps of neural activity during rhythmical activity and formation of static bumps during arrest and posture. Besides linking cell types, structure and function, we propose our approach as a new theoretical framework for motor control.

## Main

Rhythmic movements, such as walking and swimming, are orchestrated by motor networks within the spinal cord, known as central pattern generators (CPGs). These CPGs comprise a diverse array of neurons with distinct genetic profiles. Over the years, genetic tools have been instrumental in classifying neuronal cell types^2,4,6,7^, motivated by the assumption that genetic identity is directly related to specific roles in motor function^1^. While certain genetically defined neurons have been associated with fundamental roles such as rhythm generation in some species^8,9^ or with left-right and flexor-extensor alternation^10,11^, most neuronal cell types rarely exhibit a single, well-defined function^1,7,12–14^. These perplexing results imply that genetic identity alone cannot explain how connectivity patterns emerge and constrain neuronal activity. Although connections to motoneurons, including some reflex arcs, have been mapped^15^, these represent only isolated parts of the surrounding circuit, rather than the CPG network itself — whose connectivity remains largely uncharted. These gaps in understanding have led researchers to propose modular half-center models with cell-type connectivity constrained by hypothesized rules. However, this approach has limitations. First, rhythm generation is assumed to rely on pacemaker cells, whose necessity remains unsupported by experiments; second, modular models lack flexibility, as rhythmic frequency is inherently tied to amplitude, and mechanisms for slowing down and stopping while keeping the pose are unexplained. Here, we propose a solution to these open questions through principles that link cell types to spatial features and network structure to dynamics.

### Space as a principle for motor control

A key feature of spinal circuits is the generation of rhythmic activity. While pacemaker neurons have been suggested as a potential mechanism^12–14^, there is limited evidence supporting their necessity, suggesting that rhythmicity may instead arise from network-level interactions^16^. Spinal population recordings during rhythmic activity demonstrate that all phases of the cycle are represented nearly equally, a phenomenon known as rotational dynamics^17–20^. To date, only one model^17^ has successfully explained these dynamics using a randomly connected network. This raises an intriguing question: How do rhythms and rotational dynamics emerge from such simple networks without the need for specialized pacemaker cells? The solution may lie in understanding how networks can generate rhythms through their spatial organization^21–23^.

In the spinal cord of many species, a clear feature of genetically identified cell types is their exquisite spatial organization^1,4,7^. They show transverse segregation and various projection patterns in the rostrocaudal (RC), mediolateral (ML), and dorsoventral **(DV) directions. For example, in mice, a subset of excitatory neurons (V2a/Vsx2) are located in intermediate laminae along the** DV axis from where they exhibit strong projection biases in the ventral and caudal directions (**Fig. 1A-C**). Another cell type, the inhibitory interneuron, V1, is known to localize ventrolaterally and project primarily rostrally^24^. Together, the diversity of cell types forms a projection pattern with excitatory and inhibitory neurons that cover both longitudinal and transverse directions with dissimilar projection extensions^25,26^ (**Fig. S1**, and **Video S1**).

**Fig. 1.**
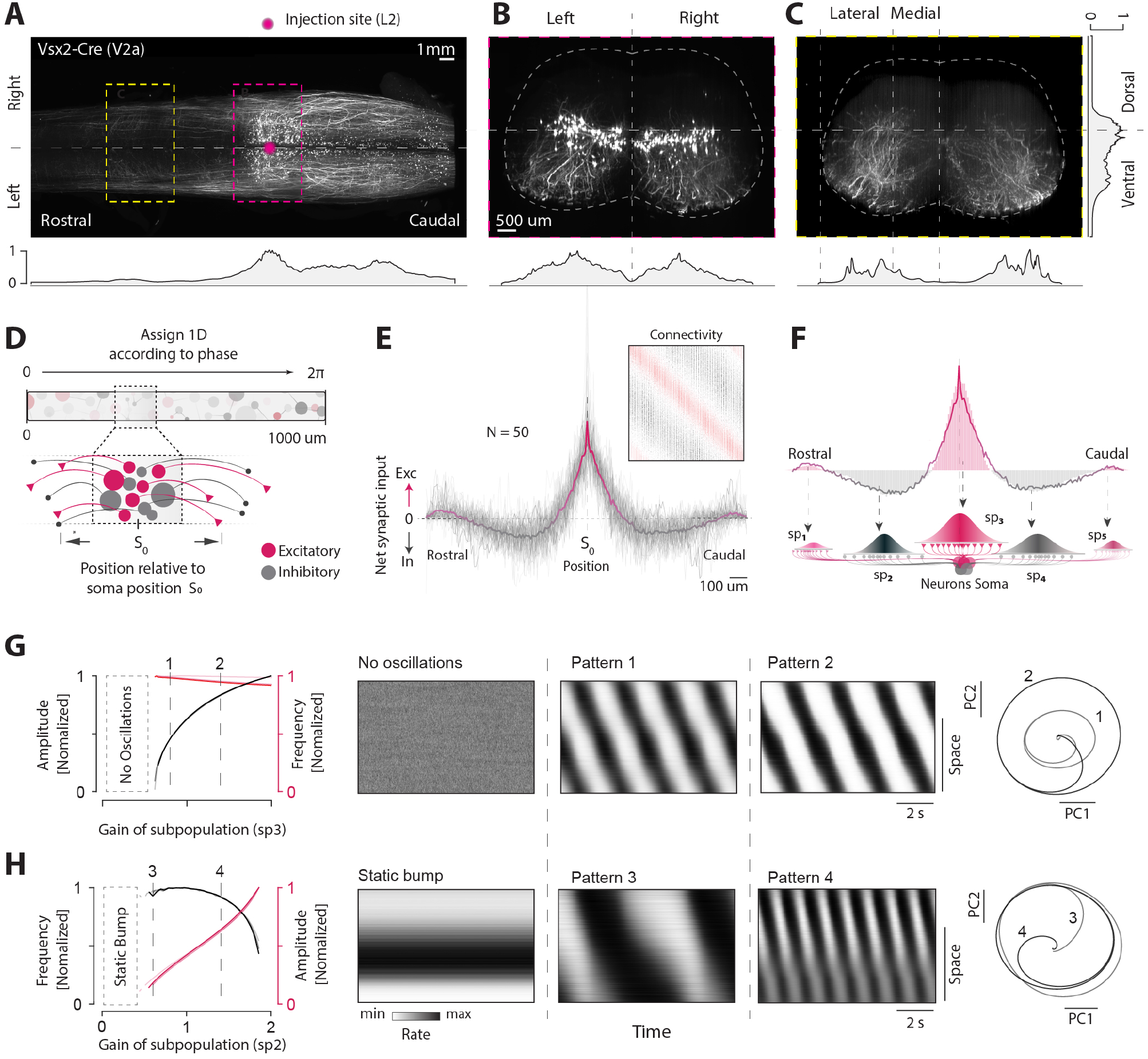
Spinal projections could form a Mexican hat network, which can generate rhythms and be controlled. **A**, Projection of a subset of the V2a excitatory neurons (Vsx2) has asymmetric bias towards rostral direction, revealed by a fluorescent reporter (AAV point injection, magenta) in the mouse lumbar spinal cord (dorsal view). **B**, Transverse section from A (magenta) shows somata and their axonal projections bias in ventral directions. **C**, Few axons also ascend (yellow highlight in A). **D**, Rhythmic model network of the cord has excitatory and inhibitory projections (red/grey) from a given location, *S*_0_. **E**, Placing the neurons according to sequence reveals a slightly asymmetric “Mexican hat”, also visible in the connectivity matrix (inset). **F**, Populating the individual lobes with projection distributions from subpopulations together comprise the net profile. **G-H**, The network generates rhythms with almost independent control of amplitude (G) and frequency (H) using gain modulation of sub-populations (left). Patterns of firing rate activity (middle) for three values of gain (indicated, left). Gain modulation of local excitation (sp3), primarily changes amplitude (black line, left), visible as larger radius in rotational dynamics (PC1-2, right). Gain modulation of rostral inhibition (sp2) primarily changes frequency (red line, left) with only small affect on amplitude, i.e. similar radius of rotation in PC-space (H, middle and right). Contrasts in A-C are individually enhanced. Data in A-C adapted from^5^.

Considering these observations, we hypothesize that spatial organization alone could serve as the foundation for motor functions.

We investigate this idea in model networks that have previously been shown to generate rhythms with rotational dynamics^17^. We assign locations to neurons by ordering them according to their sequential activity along the longitudinal axis of the spinal cord (**Fig. 1D**). The assumption is that neurons with similar phases have spatial proximity to minimize the wiring cost (**Fig. S2**). The main components of the network are the excitatory (E) and inhibitory (I) neurons, which make synapses at various targets along the line from a starting point (*S*_0_). When inspecting the net synaptic output from *S*_0_, a striking picture emerges: Local excitation and long-range inhibition form an asymmetric profile resembling a “Mexican Hat” (**Fig. 1E** and **Fig. S2**). This is also apparent in the network connectivity matrix, where local excitation and longer-range inhibition are visible as diagonal bands (inset, **Fig. 1E**). We define the net synaptic input along the longitudinal axis arising from *S*_0_ as the “projectome”. A “Mexican hat”-like projectome arises when excitatory and inhibitory projection lengths differ^21,22^ and this can easily emerge since multiple neuronal subpopulations have distinct projection patterns, as observed in spinal cord cell types (**Fig. 1A**).

To investigate the functional advantages of this projectome, we construct networks of five subpopulations (sp1-5) (**Fig. 1F, Fig. S2**). Each cell type has a Gaussian probability to form synapses with a mean projection length (*µ* ^*p*^) and width (*σ* ^*p*^). Together, the populations form the mexican hat projectome (sp1-5, **Fig. 1F**). This is the spatial principle of our theory. To cultivate this principle, we probe the network dynamics using a rate-based formalism (Supplementary Methods). All networks produce rhythmic activity in response to constant input, regardless of their size (**Fig. S2**). Next, we explore the potential roles of specific cell types by changing their gain. In particular, we find that individual populations modulate distinct rhythmic properties. For example, amplitude can be selectively increased with minimal effect on frequency by enhancing local excitation in the central lobe (sp3, **Fig. 1G**). In contrast, modulating the gain of ascending inhibitory projections (sp2) allows control of the frequency without altering amplitude (**Fig. 1H**). Interestingly, rhythmic activity can be slowed or stopped, while preserving static, localized neural activity. Similar effects are observed with other modulations of the projectome and its spatial asymmetry (**Fig. S3** and **Video S2**).

These phenomena can be explained by an asymmetry of the projectome. Symmetric network topologies, characterized by localized excitation and global inhibition, are known to support stable “bumps” of activity^23^, analogous to ring attractor dynamics in Drosophila^27^. However, when asymmetry is introduced into the projection profile, these activity bumps can transform into traveling waves^22,23,28,29^. Thus, key functional dynamics, such as rhythms, rotations, traveling waves, and slow-downs/halts, emerge from, and are modulated by, a set of cell types with asymmetric projections. We proposed this as the foundational principle of the descending control of spinal network activity.

### Constructing the mouse spinal cord

As demonstrated above, placing cell types along a linear axis can produce wave dynamics similar to those observed in undulatory species^32^. However, it has been widely assumed that control of tetrapod locomotion is fundamentally distinct due to the presence of multiple flexor-extensor muscles of the limbs in their corresponding cervical and lumbar circuits, and diversity of gaits. However, this distinction has not been rigorously tested, primarily because of experimental limitations. Given that a significant proportion of vertebrate cell types are conserved throughout evolution^1,15^, we propose that the same network principles apply broadly, allowing us to model limbed locomotion using our framework. Hence, we model the mouse spinal cord.

First, we need to determine the spatial arrangement of the cell types within the mouse spinal cord. For this, we use a combination of single nucleus **RNA** sequencing and spatial transcriptomics in a transverse section of the mouse lumbar spinal cord, previously obtained^5^ (**Fig. 2A**). According to a harmonized cell atlas^31^, we identify the locations of 69 neuronal populations (28 I, 38 E, 3 motoneurons) that span the entire DV axis. We aggregate populations of the same developmental origins^31^ and refer to these groups as our spinal cell types (**Fig. S4A**). This gives a quantitative distribution of cell types in a cross section of the lumbar cord, which resembles that established through other methods, e.g. Allen mouse spinal cord Atlas (**Fig. 2B, S4B**). We make the simplifying assumption that these distributions are preserved in the longitudinal direction and extend the map (**Fig. 2C, Fig. S5A**). This allows projections of a given presynaptic neuron to target regions in both longitudinal and transverse directions. Since CPGs can function even after deafferentation^15^, we focus on cells in the ventral territory of the spinal cord and do not include dorsal horn interneurons or sensory feedback. In addition to motoneurons (*Mn*), we include eight cell types, excitatory 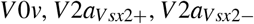 and *V* 3, and inhibitory *dI*6, *V* 0*d, V* 1 and *V* 2*b* (color coded, **Fig. 2C**). We model the T13 to S4 segments and distribute hindlimb motoneuron pools according to their identity and known segmental locations^33^ (**Fig. 2D**).

**Fig. 2.**
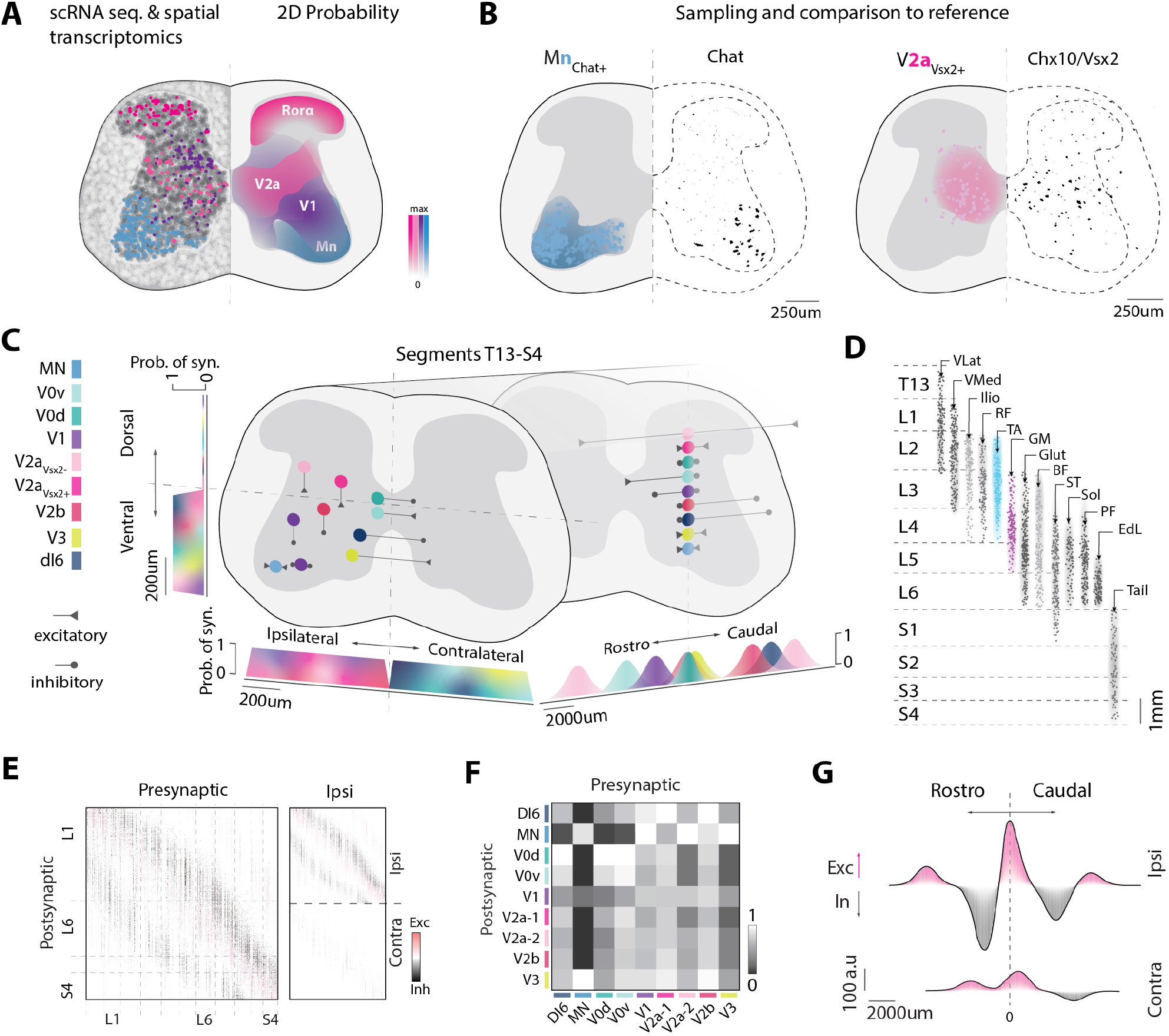
Motor function associated with spinal cell types through spatial segregation. **A**, Cell types distributions are identified using scRNA sequencing and mapped in transverse plane by spatial transcriptomics. **B**, Comparison of two types, Mn (blue) and V2a_Vsx2+_ (magenta), sampled from A, with the empirical maps of their molecular markers, ChAT and Chx10/Vsx2, respectively^30^. **C**, Nine cells types are included in a model of the lumbo-sacral region of the cord (segments T13-S4). Cell type specific projection patterns are modelled as independent Gaussians along the RC, ML, and DV axes. Each Gaussian defines a probability of forming a synapse with any target neuron falling in the distribution. **D**, Segmental hindlimb motoneurons pools distributions as assigned in the model. **E**, Left: Pairwise connectivity matrix, **W**. Right: Connectivity matrix decomposed in its ipsilateral and contralateral components . **F**, Cell type connectivity matrix (rectified and normalized). **G**, Network projectome decomposed in its ipsilateral and contralateral components. Data in A^5,31^ and B^30^ reanalyzed with permission

Next, we establish network connectivity employing similar projection rules as previously introduced (**Fig. S5A-B**). We assign spatial probabilities of projection over the three axes of space (i.e. RC, ML, and DV), which are adjusted for each cell type according to empirical observations^3,4^ (**Fig. 2C, Table S1**). For instance, V2a have both local and long-range ipsilateral projections, V0, dI6 and V3 are primarily commissural and V1 and V2b have mainly ipsilateral ascending and descending directions, respectively. A differential expression analysis of marker genes for long-range and local projections^2^ corroborates most of these choices (**Fig. S4C**). Synapses between pairs of neurons are sampled from the projection distributions of the presynaptic partner (**Fig. S5B-C**). These assumptions allow inferring the connectivity matrix (**Fig. 2E**), the connection probability between cell types (**Fig. 2F**) as well as the contra and ipsilateral contributions to the projectome, which strikingly resembles the theoretical mexican-hat (**Fig. 2G, Fig. S5C-D**, and **Video S3**).

Finally, we ask whether our spatial model constitutes a valid representation^34^ of the mouse lumbar cord. The relative fraction of cell types closely mirrors the cell counts reported for the mouse^5,31^ and the L2-L5 region exhibits a higher overall fraction of cells, reflecting lumbar enlargement (**Fig. S6A-E**). Simulated axonal projections resemble pan-neuronal axonal tracings (**Fig. S6F**) and the distributions of the premotor interneurons of a sample pair of flexor / extension muscles (Tibialis anterior and Gastrocnemius medialis) are spatially intermingled in the transverse plane but differ longitudinally, aligning with the established experimental retrograde labeling^35^ (**Fig. S7A-B)**. Hence, the model captures essential structural features of the mouse spinal cord.

We have constructed a model of the lumbar CPG that can serve both as a robust foundation for validating the approach against experimental observations and as a prediction of future experiments. We can now test whether functional neuronal activity emerges as a consequence of this network architecture.

### Emergent locomotor activity

Our objective is to associate the activity of the network with its structure. For that, we again employ the rate-based formalism to investigate the dynamics of our model^17^. Fortunately, this formalism results in a set of equations, where the connectivity matrix (**W, Fig. 3A**), provides a link between structure and function through its eigenvalues (**Fig. S8A**). Specifically, if the largest eigenvalue (dominant mode) exceeds the stability threshold and possesses an imaginary component, the network exhibits oscillatory behavior. Our model has two prominent eigenvalues associated with rhythmic activities (**Fig. 3B**), termed mode 1 and mode 2. Mode 1 is associated with four “bumps” of activity, alternating between the right and left hemicords, which we interpret as walking (**Fig. 3B, Video S4**). In contrast, mode 2 corresponds to synchrony between these “bumps”, representing a bounding mode. Interestingly, the structure of the dominant modes appears to be insensitive to the specific transverse location of the cells, and to cell-to-cell type connectivity (**Fig. S8B-C**), suggesting that they emerge as a consequence of the longitudinal projectome alone.

**Fig. 3.**
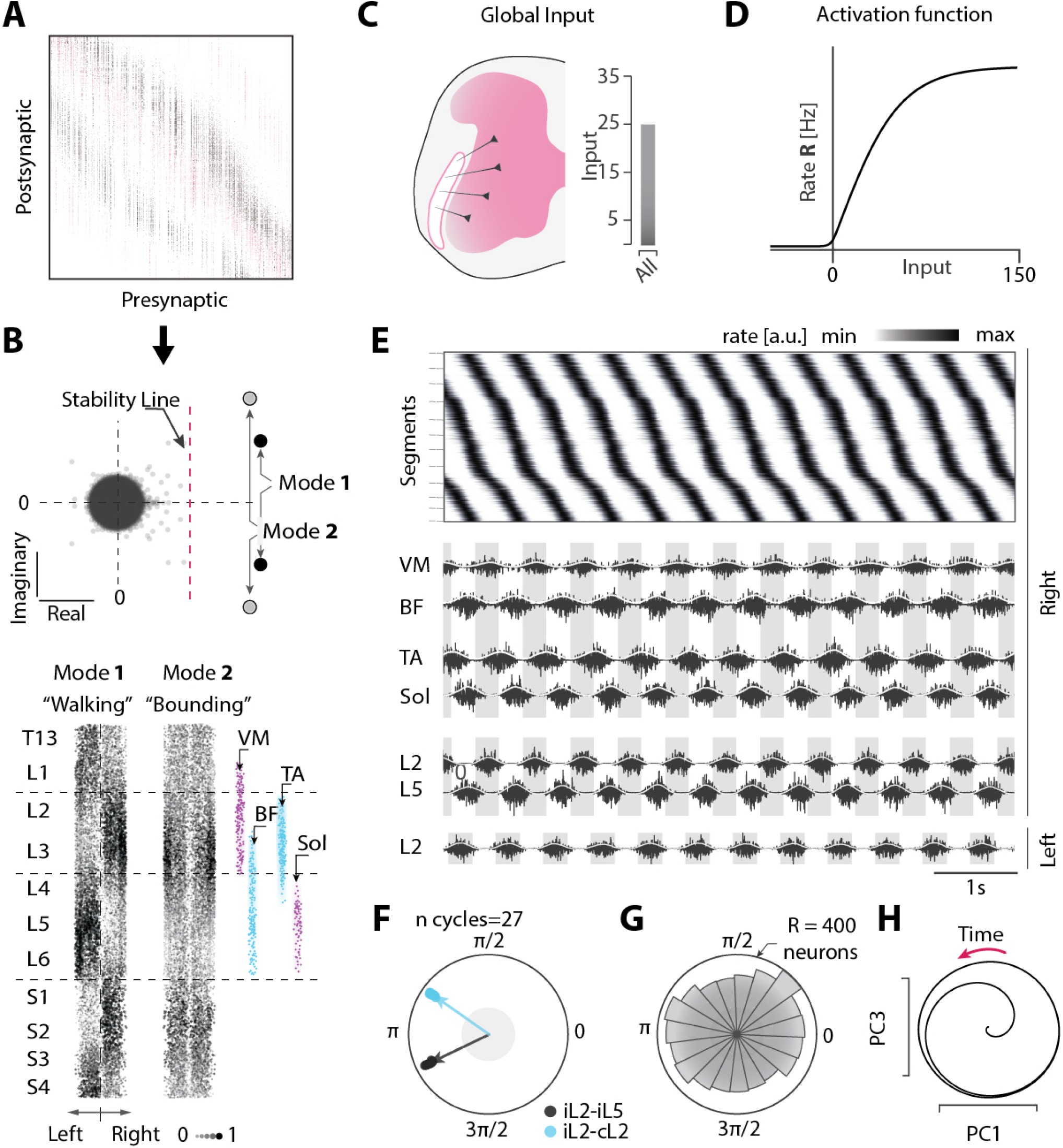
Mouse locomotion: Emergent spatio-temporal dynamics of the model. **A**, The connectivity matrix W of the network. **B**. Top: The eigenspectrum of W has two dominant eigenvalues, representing two modes, above stability line and with imaginary parts, hence oscillatory. Bottom: The corresponding firing activity (illustrated as size of dots) of neurons in space of the two modes, which represents either right-left alternation (Walking) or synchrony (Bounding). Selected flexor (blue) and extensor (magenta) antagonistic pairs (Vastus Medialis VM, Biceps Femoris BF and Tibialis Anterior TA - Soleus Sol) of motoneuron pools are indicated. **C**, Global input to the network. **D** Non-linear activation function (composite hyperbolic tangent) (**E**) Emergent dynamics. Top: Spatial representation of firing rates. Mode 1 (walking) is dominant and has “bumps” of activity traveling across segments in caudal direction (Right hemicord). Middle: Resulting muscular activity for selected pairs of muscles. Bottom: Segmental activity. **F**, Polar representation of right/left L2 phase (blue) and L2-L5 phase (black). **G**, Phase distribution of all neurons, indicates rotational dynamics. **H**, population dynamics projected onto the first two PC, displays rotation.

For comparison with the experimental data, we focus on two pairs of antagonistic motoneuron pools^33^. These motoneurons are activated by local premotor interneurons that transmit the spatial “bumps” of activity. The structure of the bumps approximates the extent and alternation of the groups of antagonist muscles (right, **Fig. 3B, Fig. S8D**). When constant input activates all neurons in the network (**Fig. 3C-D**), mode 1 naturally becomes dominant and manifests itself as periodic waves traveling rostro-caudally (**Fig. 3E**). This propagation leads to an alternating muscular activity between pairs of antagonistic muscles between sides of the cord, as well as between segments traditionally recognized as extensor and flexor centers (i.e. L2 and L5) (**Fig. 3E-F**). Although changing the position of motoneuron pools along the rostro-caudal axis preserves the dominant mode structure, it significantly alters motor output (**Fig. S8D-E**) highlighting the critical necessity of their organization to achieve functional outcomes. The population exhibits all phases and shows rotational dynamics in PCA space (**Fig. 3G-H**).

In summary, the projectome that captures the projection patterns found in the mouse spinal cord can not only generate rhythm, rotation, and patterns, but also the two main types of gait walking and bounding. Furthermore, an unbiased exploration of the parameter space of the projectome indicates that while the parameterization of the projectome is not unique, it is necessary to produce rhythmic activity. Taken together, all these elements point to spatial projection patterns as the key mechanism driving rhythmogenesis and pattern formation within the spinal cord (**Fig. S9-S11**).

### Descending control of locomotion

We next ask whether descending excitation and modulation reproduce the control of locomotion, as observed experimentally. In the rate-based framework, the firing rate of each neuron is a smooth function where the slope is the gain (**Fig. 4A**). The gain and input can be modulated by e.g. descending fibers. We introduce two descending inputs: one glutamatergic from the caudal medulla (LPGi^36^) that excites cells in intermediate laminae, and fibers from the raphe obscurus that release serotonin, which causes gain modulation in target cells in the ventrolateral spinal cord (**Fig. 4B, Fig. S11**, and **Video S4)**.

**Fig. 4.**
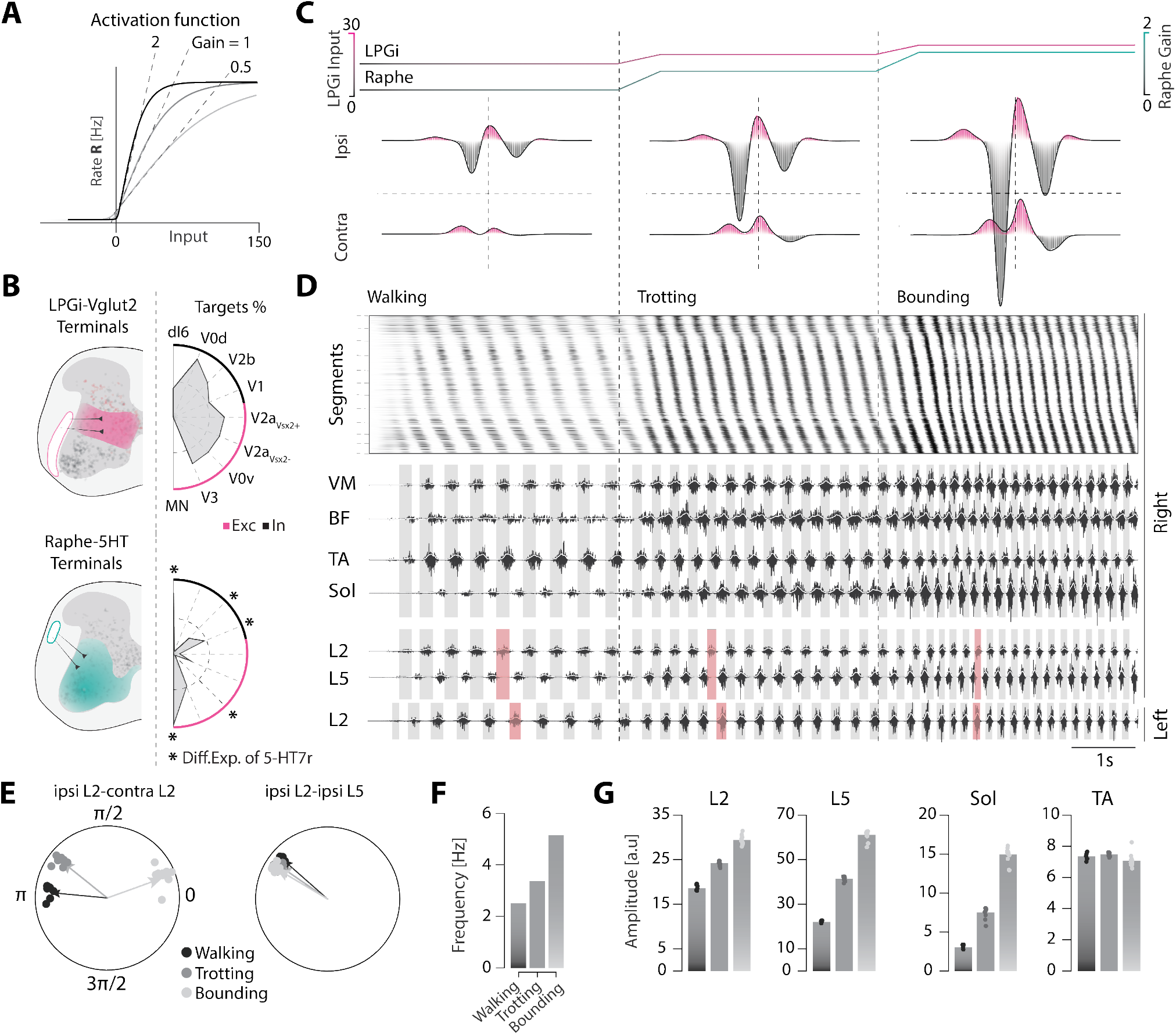
Descending control of lomotor activity by modulation of gain and projectome. **A**, Modulation of gain (slope) of the neuronal response function in the model (3 levels shown). **B**, Top: Location of the terminals of the descending glutamatergic fibers (from LPGi) and targeted area (left) and cell types (right). Bottom: Location of serotonergic fibers terminals and targeted cell types. Cell types differentially expression 5-HT7 receptors are noted with a star.**C**, Top: the descending input (increasing glutamatergic drive from LPGi, pink) and the gain of the cells receiving serotonin (increasing , green). Bottom: the corresponding effective projectome adjusted for the modulated gain. Symmetry and amplitude of the projectome change with the modulation. **D**,Left: Low frequency (walking) shows traveling waves. Middle: increase in frequency (Trotting), with same sequential muscular activity. Right: Highest frequency, with a switch in gait from left-right alternation to synchrony . The phase right/left L2 are now in phase(bottom). **G**, Polar respresentation of right/left L2 phase and L2-L5 phase. **F**, Bar graph of locomotor frequency during walking, trotting and bounding. **G**, The corresponding nerve output (L2 and L5) and muscle amplitudes of these behaviors.

We simulate network behavior under gradually increasing glutamatergic activation (from LPGi) and serotonin levels (top, **Fig. 4C**). Serotonin selectively affects populations close to the terminals of raphe obscurus, modifying the projectome in a distinct manner (bottom, **Fig. 4C**). These adjustments lead to changes in motor behavior. At low levels of serotonin, the ipsilateral projectome shows minimal asymmetry, while the contralateral contribution seems to favor excitation at long ranges, producing a slow left-right alternating pattern that mimics walking. As the 5-HT input increases, the ipsilateral projectome becomes increasingly asymmetric while contralateral contributions shift in favor of local excitation and longer range inhibition, giving rise to faster locomotor patterns such as trotting and bounding (**Fig. 4D-G**). This progression is also reflected in the shifting positions of the eigenvalues and relative changes in firing rates (**Fig. S11**). The specific changes in dominance of the contralateral projectome is the fundamental mechanism for gait changes as revealed by the unbiased parameter search performed previously (Supplementary Method, **Fig. S9**) and corroborated by experimental observations^37^. A global excitatory input can activate the network as effectively as specific input from LPGi. However, rearranging the cell positions in the transverse plane renders only the global input effective, while the LPGi input fails to produce rhythms (**Fig. S10**). Similar results are observed with raphe gain modulation (data not shown), indicating that the transversal position is essential for descending tracts to target the minimal group of cells required for proper function. This aligns with observations that both pharmacological activation and brainstem stimulation can induce rhythmic activity in the spinal cord.

In summary, our model of the mouse spinal cord, grounded in the principle of linking structure with function and cell types, successfully generates activity that aligns with realistic motor behaviors and recapitulates important features of descending control. The explicit incorporation of genetic cell types allows now to compare computational manipulation experiments with reported experimental findings.

### Manipulation of cell types

A common experimental strategy has been to perturb cell types and characterize the effects on activity. An example is a type of inhibitory interneurons (V1), which can be manipulated tractably using various genetic approaches. In general, silencing of V1 neurons causes a slowdown of locomotor rhythmicity^38,39^. This slowdown is counterintuitive and has been difficult to explain. Furthermore, perturbations of V1 neurons appear to have other effects than frequency alone, such as an increase in the spiking and flexor-extensor duty cycle^40^. Could these and other previous experiments, such as silencing V0 be explained in our framework?

Here, we first perform a “silencing” of the V1 interneurons in the model. This is done selectively first by hypopolarization and then ablation (**Fig. 5A-C**). The result is a slowdown, in agreement with the experiments. In addition, the burst amplitude increases, whereas the left/right and flexor/extensor phase relations are unaffected (purple **Fig. 5B-C**). Although this agreement is reassuring, it does not itself explain the mechanism. We suggest the following explanation, which is rooted in the spatial principle: In the intact situation, the ascending inhibition (*sp*_2_, **Fig. 1F**) prevents the spread of local excitation in the rostral direction. This effect, combined with an excitatory projection bias towards the tail, causes the bump of activity to propagate caudally. When decreasing or eliminating the ascending inhibition (*sp*_2_), the local excitation bump will remain for a longer time, which effectively increases the descending inhibition and therefore causes a slowdown in caudal propagation. The reverse is also true: When the ascending inhibition is strengthened, the bump will move faster in the caudal direction, as seen when increasing the gain of *sp*_2_ (**Fig. 1H**). A similar argument also applies for the descending inhibition (*sp*_4_), which is provided by V2b cells^41^ in the mammalian spinal cord. In reality, this is more complicated since these fibers are less numerous than V1 cells^11^, and their respective ratio ensures an asymmetry in longitudinal inhibition. In addition, the amplitude of the motor output is generally increased due to the imbalance of the E / I ratio when silencing V1 or V2b. Silencing both V1 and V2b will result in an uncontrolled fast spread of excitation, resulting in a collapse of the flexor / extension alternation^11^. A similar effect is observed in the model upon blocking of inhibition (data not shown).

**Fig. 5.**
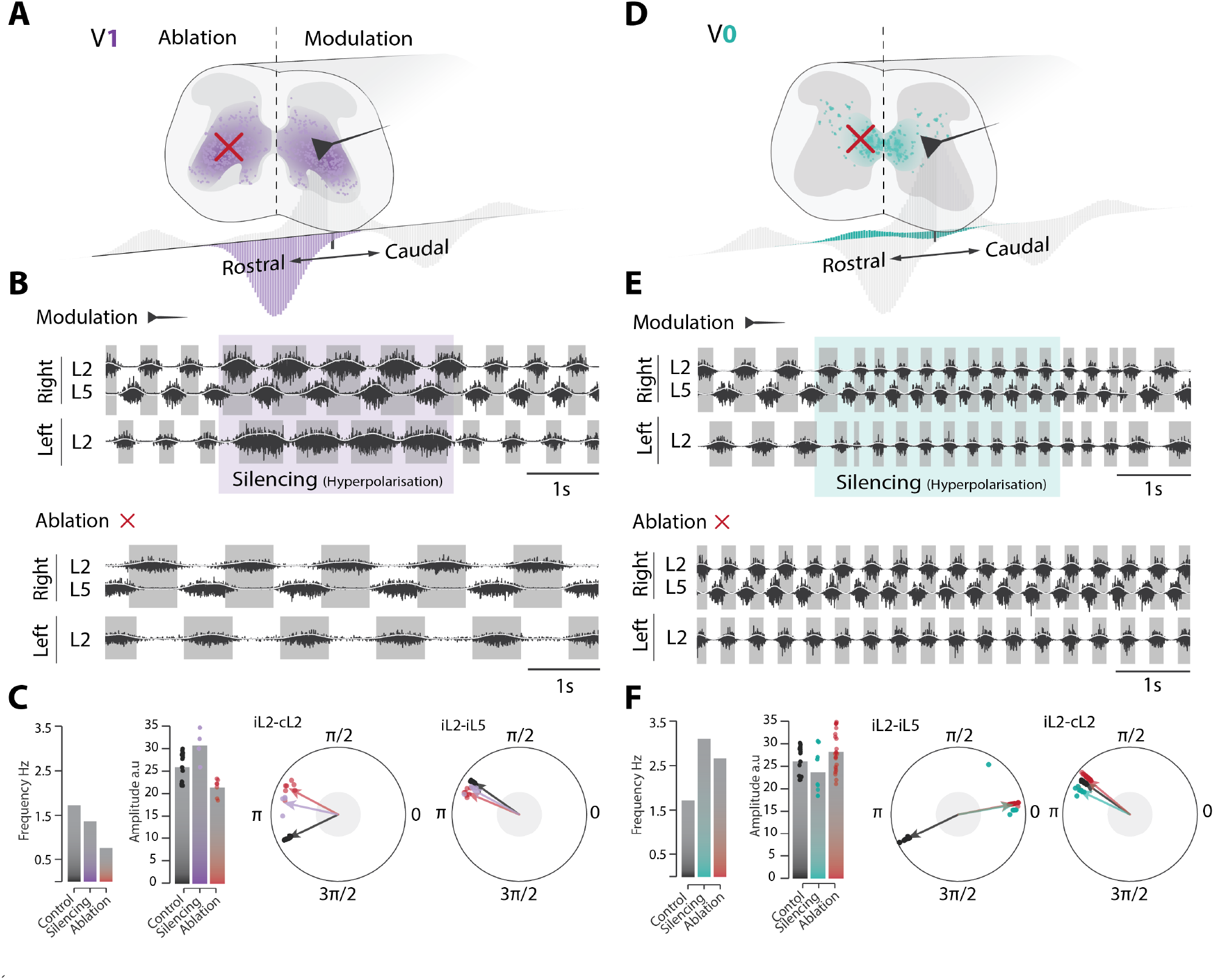
Manipulation of V1 and V0 cells in model. **A**, V1 cell type is manipulated either by hyperpolarization, imitated by a black synapse, or simply removed from the network (ablated, red cross). Only the rostral lobe of the projectome is affected (purple distribution, bottom). **B**, Slowing of motor nerve activity of spinal segments L2 and L5 as a concequence of V1-silencing (top, purple area), and after V1-ablation bottom. **C**, Mean frequency decreases with silencing and ablation, while the amplitude is increase (silencing) or decreased (ablation, bar graphs, left). Phase relations of right/left (L2) and ipsilateral (L2-L5) are large unaffected (polar plots, black: control, purple: silencing, red: ablation). **D**, V0 manipulation by silencing or ablation. Contralateral contributions to the spinal pojectome is now affected (cyan, bottom). **E**, Motor nerve activity before and during V0-silencing has two consequences; a speed up and a right/left phase shift (cyan area, top). Ablation has similar effects (bottom). **F**, Mean frequency increases during V0 silencing (cyan) and ablation (red) compared with control (black). The amplitudes are largely unaffected (right bar graphs). The right/left phase of L2 switch from alternation to synchrony (cf. black, cyan and red, left polar plot) while the phase of ipsilateral L2-L5 remains unaffected (right polar plot).

Another experimental manipulation that has been performed is targeting the V0 interneurons using genetic ablation. This caused quadrupedal hopping at all locomotion frequencies^37^. We test the effect in our model by first hyperpolarizing V0, which alters the commissural projectome^26^ (cyan **Fig. 5D**). Hyperpolarization causes both an increase in locomotor frequency and a right-left L2 synchronization, that is, a hopping (cyan area, top **Fig. 5E**). Complete ablation of V0 cells has similar effects (bottom). The ipsilateral phase of L2-L5 and muscle amplitudes are largely unaffected, while the right-left coordination changes (**Fig. 5F**).

In conclusion, we demonstrate that previous experimental observations can be reproduced and explained by mechanistic principles. The function arises out of dynamics, dynamics arises out of the network, and the network is built out of cell types with spatial segregation and projection biases. The framework also turn out to be remarkable robust to pertubation, and does not require parameter optimization for producing sensible motor activity (**Fig. S8-S12**).

## Discussion

Although cell types are often assumed to be related to spinal motor function^1,4,42^, understanding how they are linked has been absent from the literature. An important clue is that many molecular markers are associated with axon guidance, cell adhesion, or general spatial organization^31^. Here, we propose that motor functions emerge out of spatial structure, where the location of cell types together with their longitudinal projections constitute the principle of organization. Asymmetric projections with local excitation and long-range inhibition are essential for rhythm generation and sequential activation of motoneurons. The link between cell type and function is established through the dynamics of the network, which is shaped by a combination of the projectome and spatial location.

We test our approach in models, where the spatial distribution of cell types is extracted empirically using single cell sequencing combined with spatial transcriptomics and where the cell type projectome composition is gleaned from the literature (**Fig. 2**). We find that dissimilar projection lengths readily give rise to an asymmetric “Mexican hat” organization, i.e. local excitation and long-range inhibition. The asymmetry causes an emergent bump of local activity to propagate in the caudal direction (**Fig. 3**). Since the shape and size of the projectome determine properties of the activity, e.g. frequency and amplitude (**Fig. 1**), modulation of individual cell types that make up the projectome constitutes a convenient means of descending control. Hence, modulation of cell types with well-characterized effects, e.g. V0 and V1, resulted in behaviors in agreement with experimental reports (**Fig. 5**). In particular, our model is capable of generating appropriate behavior without requiring fine adjustment of particular synaptic connectivity weights or cellular mechanisms such as in pacemaker properties (**Fig. S8-12**).

This approach stands in contrast to conventional CPG models where the network is constrained by posited rules, e.g., modules with reciprocal inhibition, feedforward excitation, and where rhythms are generated by cellular pacemaker properties^15,42^. This modular organization, inspired by the half-center network^15,42^, has difficulty explaining the gradual slowdown and stopping of movement. This is due to both the autonomous nature of rhythmogenic cells and the abrupt transition of activity between antagonistic modules with recurrent unbalanced excitation^43^. When the agonist modules are active, the antagonistic modules are kept silent by reciprocal inhibition, and vice versa. Hence, all cells have the same or opposite phase, and this is in disagreement with the continuous progression of cyclic activity, that is, the rotational dynamics of the population, observed throughout the neuronal population^17–20^. Instead, we propose a spatial organization in which the adjustment of the excitatory and inhibitory lobes of the projectome allows full control of all aspects of rhythmic activity, including slowing and stopping in any phase of the cycle (**Fig. 1, Fig. S2**).

Our approach predicts traveling waves in the spinal cord during tetrapod locomotion. This idea is not new and has been proposed and demonstrated in various species^24,44–47^. The propagation of waves in limbed vertebrates has also been demonstrated, for example in frogs^48^, salamanders^49^, and newborn rodents^50,51^, which could also explain a wave within motoneuron pools^51–53^. Systemic block of inhibition leads to a fast seizure-like wavefront that effectively synchronizes nerve activity^54^. Surface field potentials show sinusoidal waves in cats under fictive rhythmic scratching^55,56^. However, another report was unable to support such a wave among interneuron activity^57^, suggesting that the issue could be more complicated and even specific to cell types if it is a propagating wave. This issue remains to be investigated further in future studies.

Ultimately, our approach, with its clearly defined physical structure, lays the foundations for investigating motor diseases that affect the spinal cord’s integrity, such as spinal cord injuries. By incorporating the transcriptome of each cell, it opens exciting possibilities for studying how to restore motor functions through genetic targeting and regrowth of specific cell population projections within the cord.

## Acknowledgements

This work was supported by The Lundbeck Foundation (no. R366-2021-233, R.W.B.), Novonordisk foundation (no. NNF23OC0082192, R.W.B), and The Swiss National Science Foundation (no. SNF Postdoc.Mobility P500PB_206824, S.K.). We would like to thank Prof. Grégoire Courtine for generously sharing the Vsx2 tracing data. We extend our gratitude to Prof. Claudia Kathe for her help and technical expertise.

## Author contributions

R.W.B., S.K. A.W. conceived the original idea. G.H., G.L., and M.C.A.B. collected the data. S.K. and A.W. performed the model simulations. S.K. A.W. and R.W.B. designed, and developed the theory. G.H., T.T., R.J.S., S.D.L., M.C.A.B., G.L., S.K., R.W.B., analyzed the experimental data. S.K., R.W.B. wrote the manuscript.

## Competing interests

The authors declare no competing interests.

## Notes

### Competing Interest Statement

The authors have declared no competing interest.

### Summary of Updates

Text improved and author added. Figures have been updated.

