## Supplementary Information for "Spatial and network principles behind neural generation of locomotion"

### Spinal Injections, Tissue Clearing, and Imaging

The data and methods related to Figure 1A-C were provided and reanalyzed with permission from a previous publication[1]. The images in Fig. 1A-C were subjected to individual contrast enhancement. The data of Fig S1 and Video S1 (**Fig. S1 and Video S1**) were done using wild-type mouse in the following steps.

**AAV Injections** Wild-type mice (C57BL/6) were injected with an adeno-associated virus (Addgene viral prep 50465-AAV9) that carried the fluorescent reporter EGFP, driven by a neuron-specific promoter (hSyn), into the lumbar spinal cord. The animals were sacrificed 4 weeks later via a lethal dose of anesthesia followed by a transcardiac perfusion with 20 mL of PBS followed by 20 mL of 4% PFA solution. The tissue was post-fixed overnight at 4°C and then washed with PBS azide and kept at 4°C until the start of the tissue clearing and immunolabeling protocol.

**Tissue Clearing and immuno labeling** To enable imaging of the entire central nervous system (CNS), tissue clearing and antibody labeling protocols were optimized to enhance the visualization of fluorescent signals throughout the CNS. The tissues were processed using a modified iDISCO+ protocol[2]. The samples were pretreated with 25% quadrol (v / v) in water for 2 days at 37°C, followed by 5% ammonium hydroxide (v/v) in water for 1 day at 37°C. Residual chemicals were removed through multiple washes in PBS containing 0.02% sodium azide at room temperature (RT). Dehydration was performed using a series of graded methanol (20%, 40%, 60%, 80%, 100% v/v in water) for 1 hour each at RT, with two additional steps of 100% methanol. Lipids were removed by incubating the tissues in 2:1 dichloromethane (DCM) / methanol overnight (O / N) in RT, followed by multiple washes in 100% methanol. Samples were bleached with 5% hydrogen peroxide ($H_2O_2$) in methanol O/N at 4°C, rehydrated stepwise (60%, 40%, 20% methanol solutions in water for 1 hour each at RT), and washed in PBS for 1 hour at RT.

Permeabilization was performed in a solution containing PBS, 0.02% sodium azide, 0.2% Triton X-100, 10 $\mu$g/mL heparin, 2.3% glycine, and 20% dimethyl sulfoxide (DMSO) for 1 day at 37°C, followed by blocking in PBS with 0.02% sodium azide, 0.2% Triton X-100, 10 $\mu$g/mL heparin, and 3% donkey serum for 1 day at 37°C. Samples were incubated with primary antibodies (GFP: GFP-1020, Aves; RFP: 600-401-379, Rockland) at dilutions of 1:3000 (GFP) and 1:1000 (RFP) in antibody incubation solution (PBS containing 0.02% sodium azide, 10% w/v CHAPS, 1% w/v cyclodextrin, 1% w/v glycine, 10% v/v DMSO, and 2% v/v Triton X-100) for 10 days at 37°. After extensive washing in PTwH solution (PBS with 0.02% sodium azide, 0.2% Tween-20, and 10 $\mu$g/mL heparin) at RT and a final extended wash for 1 day, tissues were incubated with secondary antibodies (647 donkey anti-rabbit: 711-605-152, Jackson Immuno; 647 donkey anti-chicken: A78952, Thermo Fisher) at 1:1000 dilution (diluted in PBS with 0.02% sodium azide, 0.2% Triton X-100, 3% donkey serum, and 10 µg/mL heparin) for 7 days at 37°C. Following an overnight wash in PTwH at RT, samples were re-dehydrated through a graded methanol series (20%, 40%, 60%, 80%, 100% v/v in water) for 1 hour each at RT, with two additional 100% methanol steps. Lipid removal was completed by incubating tissues in 66% DCM/33% methanol O/N at RT, followed by two washes in 100% DCM for 30–45 minutes each. Finally, tissues were cleared in ethyl cinnamate (ECi) for long-term storage and imaging.

**Imaging** Acquisitions were performed with a 3i Cleared tissue lightsheet microscope and an Ultramicroscope Blaze (Miltenyi Biotec) lightsheet microscope. The microscope was installed on an air table to reduce vibration artifacts.

**3i** Two 5x illumination objectives (0.14NA) were used to generate dual side illumination of the sample, together with a 1x detection objective (Carl Ziess, 1.0x/0.25NA, WD 18mm) and a 3.25 axio zoom (2 $\mu$m/pixel). The images were acquired with a Hamamatsu ORCA-Fusion BT C15440 sCMOS camera with pixel number 2304×2304, pixel size 6.5$\mu$m, 95% QE using Slidebook software. The laser line used to capture either the RFP or GFP signal (which was enhanced with an Alexa Fluor® 647 secondary antibody) was a 637 nm, 140 mW solid-state diode laser.

**Blaze** Two illumination objectives (0.0135–0.135 NA) that generate 3 bidirectional light sheets together with a 4x and 12x detection objective (MI PLAN 4x: 0.35NA, WD 16mm and MI PLAN 12x: 0.53NA, WD 10.9mm) without zoom. The images were acquired with a 5.5 Mpx sCMOS camera, pixel size 6.5µm using ImSpector software. The laser line used to capture either the RFP or GFP signal (which was enhanced with an Alexa Fluor® 647 secondary antibody) was a 639 nm, diode laser.

**Whole-spinal-cord segmentation of neuronal projections** Axonal projection segmentation was performed using a workflow inspired by the DeepTRACE pipeline. The 3D images were processed using open access software (TRAILMAP)[3] with default parameters, and another open access analysis tool for light sheet microscopy files and which is optimized for the pretrained model 3 from DeepTrace, DeepTrace). This processing generated a 3D probability map of axons, which was projected onto the longitudinal axis to quantify local, rostral, and caudal projections. The 640-nm spinal cord scans, acquired at 2 $\mu$m isotropic resolution, were divided into overlapping chunks ( 2 mm per chunk, with 10% overlap) along the rostro-caudal axis. Each chunk was segmented using the three models from the DeepTRACE pipeline. The output of the respective models was combined by selecting the highest probability value for each voxel across all models. The combined probability maps were then subjected to iterative binarization at eight thresholds ranging from 20% to 90% of the maximum intensity value. The binarized segmentations were skeletonized, and the resulting skeletons were weighted by their respective probability thresholds before being summed together. Small disconnected components (<100 voxels) were subsequently removed from the skeletonized data. Finally, the cleaned skeletons of all data chunks were merged to reconstruct the entire spinal cord skeleton, and projection density was estimated throughout the rostro-caudal extent of the specimen.

**Immunohistochemistry** All immunohistochemistry images were adapted from the Allen brain Atlas of the Mouse Spinal Cord (Allen Brain Atlas)[4].We exported images from the expression mask, inverted their color and converted them to gray scales. Some linear transformations (shearing, scaling) were applied to fit our layout.

### Models and simulations

In all simulations presented in this report, we employed a rate model formalism governed by the following differential equation:

$$\tau_i \dot{x}_i(t) = -x_i(t) + \sum_{j=1}^{N} w_{ij} F(x_j(t)) + I_i(t) + \varepsilon_i(t)$$

where

$$F(x) = \begin{cases} T \cdot \tanh\left(\frac{g(x-T)}{T}\right) + T, & \text{for } x \leq 0 \\ f_{\max} \cdot \tanh\left(\frac{g(x-T)}{f_{\max}}\right) + T, & \text{for } x > 0 \end{cases}$$

Here, the subscripts denote the neuron index. The parameter $\tau$ (set to 50 ms) represents the integration time constant, $x_i(t)$ is the input potential of neuron $i$, and $\dot{x}_i(t)$ is its time derivative. $w_{ij}$ is the synaptic weight of neuron $j$ impinging on neuron $i$. $I_i(t)$ is the external input to neuron $i$, and $\varepsilon_i(t) \sim \mathcal{N}(0, 0.4\,\text{mV})$ is a Gaussian noise term with a mean of 0 mV and standard deviation of 0.4 mV. The function $F(x)$ is a compound tanh activation function that maps input potential to firing rate, with $T$ as a threshold parameter, an intrinsic gain $g$ and $f_{\max}$ as the maximum firing rate (**Fig S2B**). This function ensures nonnegative firing rates across all neurons. We numerically integrated this equation for each neuron in the network using the Euler method with a time step of $\Delta t = 1$ ms. The output of each simulation was a matrix that represents the firing rate of every neuron over time.

#### Random Networks

100 random networks of size $N = 500$ were generated. In all networks, excitatory (inhibitory) neurons constituted $1/2$ of the $N$ units. A synapse from neuron $j$ to neuron $i$ was placed in an $N \times N$ matrix $\mathbf{W}$ (**Fig. S2**), where columns denote source neurons and rows denote target neurons, using a binomial draw over the synapse probability, $p_{\text{synapse}} = 0.1$. Synaptic weights $w_{ij}$ were sampled from a Gaussian distribution $w_{ij} \sim \mathcal{N}(\mu = 1, \sigma = 0.1)$. According to Dale's principle, inhibitory synapses were multiplied with $-1$ so that the inhibitory and excitatory columns in $\mathbf{W}$ were strictly nonpositive and nonnegative, respectively. A detailed balance between excitation and inhibition[5–7] was imposed on $\mathbf{W}$ such that $\sum_{j=1}^{N} w_{ij} = 0$ for each row $i$. Finally, the diagonal of $\mathbf{W}$ was set to zero, i.e. removing self-synapses. Random oscillatory networks were assigned a 1D spatial coordinate based on a phase-space coherence rule (**Fig. S2F**). From the connectivity $\mathbf{W}$ and the coordinates, the synapse-weighted relative distances at which excitatory and inhibitory units tended to project was averaged over all (n=50) networks (**Fig. S2G**).

#### Spatial Unfolding

The Balanced Sequence Generator (BSG) model of rotational dynamics consists of a sparse network with randomly coupled excitatory and inhibitory neurons subject to local balance between excitatory and inhibitory synapses[8]. The dynamics of such networks may be predicted by linear stability analysis, where one examines the spectrum of eigenvalues belonging to the network connectivity matrix $\mathbf{W}$ (see **Mathematical Note: Spectral analysis**). The BSG model was proposed as a motor generating circuit in the spinal cord[8], as it is so far the only model capable of generating rhythms without pacemaker cells and generating rotational dynamics (**Fig. S2A-D**), i.e. exhibit sequential activity with a continuous distribution of phases (**Fig. S2D**). It also allows independent control of amplitude and frequency and explains other essential features of movement. However, the BSG model also has limitations. One limitation is that only some network instantiations have the rhythmogenic property. A second limitation is that the frequency is related to the size of the network (**Fig. S2E**). The dominant frequency decreases as the size of the network increases. Both effects are small-size effects due to some random networks having a dominant eigenvalue (i.e. the rightmost eigenvalue) above the stability line with a nonzero imaginary part (red arrows **Fig. S2A**). For larger networks, the eigenvalue distribution approaches the unitary circle[9]. Hence, the dominant mode moves close to the x-axis and the frequency decreases. In addition, large random networks exhibit a sharp transition from quiescence to chaos, effectively prohibiting the rhythmic state from emerging[10,11]. To circumvent these problems, we aim to pinpoint the features of small networks that give rise to rhythms and use them to build a more detailed model involving space and cell types, which can be directly tested in experiments. We start by unfolding the BSG model, i.e. assign spatial location to neurons in the network, to identify features that enriched the network with rhythmogenic properties.

The oscillatory networks were unfolded on a line according to their phase, i.e. assuming that correlation in phase reflects spatial proximity (**Fig. S2F-G**). Examining the relative distances at which excitatory and inhibitory units tended to project revealed an asymmetric "Mexican Hat" over the relative distances, that is, where excitation dominates locally with inhibition at longer ranges (**Fig. 1E-F** and **Fig. S2G-H**). We then adapted such an asymmetric "Mexican Hat" structure and imposed it on networks through subpopulations with distinct distance-based projection biases. This gave rise to consistent sequential activity at stable frequencies across network sizes (**Fig. S2H-J**). Although asymmetry is not sufficient for rhythm generation, it is necessary. It is possible, as we can see in a simulation (**Video S2**), to stop the propagation of a wave by adjusting the projectome where it is not symmetric in this case.

### Scaling of Networks

To investigate the dependence of network size, compared to their ability to generate rhythms, we constructed networks of varying sizes ($N = 200$, $N = 500$, $N = 1000$, $N = 2000$, $N = 5000$, and $N = 10000$). These networks had the imposed structure based on asymmetric Mexican Hat profiles (**Fig. S2A-G**). First, the $N$ neurons in each network were divided into five subpopulations: sp1, sp2, sp3, sp4, and sp5 (**Fig. S2H**). Second, regardless of type, each neuron was assigned a 1D spatial coordinate $x$, evenly spaced and drawn uniformly from $[0, L_N]$, where $L_N$ is the length of the network. This ensured equal representation of subpopulations along the line. We use probabilistic connections between the pre- and post-synaptic neurons, where the spatial location of the post-synaptic neuron, i.e. the distance between two neurons, is the input variable. To instantiate connections, a Gaussian distribution $\mathcal{N}(\mu^p, \sigma^p)$ was fit to each bump on the unfolded Mexican hat profile and assigned to each subpopulation as a projection probability distribution over relative distances, $d_{ij} = x_i - x_j \in [-L_N, L_N]$. Thus, a synapse from neuron $j$ to neuron $i$ was placed with probability

$$P_{syn}(i, j) = G_p \mathcal{N}(d_{ij} | \mu^{p(j)}, \sigma^{p(j)})$$

where $\mu^{p(j)}$ and $\sigma^{p(j)}$ are mean and standard deviation of projection distances of subpopulation $p$ neurons, respectively. $G_p$ is a scaling parameter to adjust network sparsity. Subsequently, the binary adjacency matrix $\mathbf{A}$ was built through the binomial sampling process

$$\mathbf{A}(i, j) \sim \text{Binomial}(1, P_{syn}(i, j))$$

where $\mathbf{A}(i, j) = 1$ indicates a synaptic connection between neurons $i$ and $j$. Synaptic weights were obtained by element-wise multiplication (Hadamard product) of $\mathbf{A}$ with the synaptic gain matrix $\mathbf{G}(i, j) \sim \mathcal{N}(1, 0.1)$, where the columns belonging to excitatory sp1, sp3 and sp5 and those belonging to the inhibitory sp2 and sp4 were strictly nonnegative and nonpositive, respectively. This yielded the full connectivity matrix $\mathbf{W}$ of the network as the Hadamard product of $\mathbf{G}$ and $\mathbf{A}$:

$$\mathbf{W} = \mathbf{G} \circ \mathbf{A}$$

Finally, a local excitatory / inhibitory balance was imposed on connectivity such that $\sum_{j=1}^{N} w_{ij} = 0$ for all target neurons $i$.

Based on these conditions, we built networks with the imposed Mexican hat-like structure. The connectivity matrix had diagonal bands indicating this projection structure (**Fig. S2I**). Having such a blue-print structure all the instantiations of networks both had oscillatory properties with a stable frequency (**Fig. S2J**). Next, we manipulate the gain of the individual subpopulations to see how we can control the activity and propagation of waves.

### Wave Propagation Mechanism

Classical work on spatially structured networks with homogeneous projections have demonstrated their pattern-generating properties in space[12–14]. To motivate the mechanism by which our network theory produces rhythms and patterns, consider a variant of Amari's one-dimensional field model

$$\frac{\partial u(x,t)}{\partial t} = -u + \int_{\Omega} w(x-y) F(u(y,t)) dy \tag{1}$$

where the spatial domain $\Omega = [0, 2\pi]$ is periodic, $u(x,t)$ is the average activity of neurons at position $x$ at time $t$ , $w$ the Mexican Hat coupling function and $F$ is sigmoid nonlinearity. If $w(x)$ is symmetric, meaning that $w(x) = w(-x)$, networks following (1) support the emergence of spatially periodic patterns - the neural analog of Turing patterns in reaction-diffusion systems. Such instabilities can arise in the linear regime, where small perturbations $v(x,t) = u(x,t) - u_0$ to the uniform fixed point $u_0 = \int_\Omega w(x-y)F(u_0)dy$ are governed by

$$\frac{\partial v(x,t)}{\partial t} = -v + F'(u_o)\int_\Omega w(x-y)v(y,t)dy. \quad (2)$$

We look for eigensolutions of the form $v(x,t) = e^{\lambda t}v(x)$ . Substituting into equation (2) yields the eigenvalue equation:

$$(\lambda+1)v(x) = F'(u_o)\int_\Omega w(x-y)v(y)dy. \quad (3)$$

Since $w$ is symmetric, spatially periodic perturbations $v(x) = e^{ikx}$ constitute eigenfunctions of (3) with $\lambda(k) = -1 + F'(u_o)\hat{w(k)}$. Hence, the growth and decay of Fourier modes with spatial frequency $k$ is determined by the intrinsic network gain $F'(u_0)$ and the magnitude of the kernel Fourier transform $\hat{w(k)}$.[15–17]. If, however, projections are asymmetric in space such that $w(x) \neq w(-x)$, resting state instabilities can give rise to patterns that propagate over space in time, much like flow or advection in pattern forming systems[15,18]. Such asymmetries serve as candidate mechanisms for generating spatio-temporal activity sequences in spatially distributed networks[19]. Spatially linear Mexican Hat networks can be realized from five underlying subpopulations, $sp_1, sp_2, sp_3, sp_4$ and $sp_5$, each contributing to the five distinct lobes on the projectome via their projection biases (**Fig. S3**). Varying these biases (**Fig. S3A-C**), or the subpopulation gains (**Fig. S3D**), impacts the asymmetry of the projectome in a manner which efficiently controls 1) the frequency of spatio-temporal sequences and 2) the direction of the travelling bump pattern. We conjecture that spinal locomotor networks function with this mechanism. In our theory, adjusting the effective asymmetry of Mexican Hat connectivity along the rostro-caudal spinal axis provides a parsimonious mechanism for rhythm- and pattern generation in animals. It also provides a straightforward means for the descending input to control the rhythm and motor activity. The next step is to include the spatial distribution of cell types in the transverse plane to match these to the sub-populations in the model, i.e. which cell types that play which role in the projectome. The purpose of this is to investigate a potential link between cell type and function and how that can be explained in our framework.

### Estimation of 2D Spatial cell type distributions

To determine the spatial distributions of cell-types unbiasedly, we used a combination of spatial omics data. Curated and annotated single cell / nucleus RNA sequencing and spatial trasncriptomics data sets of the lumbar spinal cord were obtained from the Gene Expression Omnibus: GSE184370 (single nucleus[1]), GSE184369 (spatial transcriptomics[1]), and the SeqSeek repository[20]. Our primary single-cell data set was the SeqSeek-labeled data, which classifies neurons into 69 genetically distinct populations: 28 inhibitory, 38 excitatory and 3 motoneuron-related (**Fig. S4A**). We used the developmental lineage labels from[20] as a coarser grouping to define our cell types for further analysis.

We deconvolved the spatial transcriptomics data with the labeled single nucleus RNA sequencing data using robust cell-type decomposition (RCTD)[21]. We applied default parameters, with the doublet mode disabled, yielding a matrix of 9755 2D spatial locations x 69 populations. This matrix assigns a weight to each population for each spatial location. After logarithmic normalizing and scaling the matrix columns to a range [0,1], we applied a cutoff at the mean weight to sharpen the spatial densities. The resulting distributions of each cell type closely aligned with their known locations as verified through other methods, e.g. immunostaining and in situ hybridization (**Fig. S4**). The distributions of the cell types of the sample are shown (left side) and compared to the hybridization in situ (right side, based on Allen mouse spinal cord Atlas) of a prominent genetic marker of the particular cell type (**Fig. S4B**). Genetic markers, which are associated with long-range rostro-caudal projections, Zfhx3 and Foxp2 (left, **Fig. S4C**) or local projections, Nfib and Neurod2 (right), seemed to agree closely with those previously reported[22].

### Mouse Network Construction and Connectivity

Now, we have established spatial distributions of the most generic cell types in the lumbar spinal cord. Next, we will match locations of neurons with the synaptic target somewhere else in the spinal cord based on probabilities. We first defined the spatial arrangement of the neurons based on single cell RNA sequencing and spatial transcriptomics. This defined the spatial probability distributions of various cell types (**Fig. S5A**). We then extend this in the longitudinal direction assuming the distribution is preserved (right). Now, we have neuronal populations distributed in the spinal cord. The next step is to connect these neuronal population under the assumption that cells connect probabilistically based on bias in projection which is different for each cell type (**Fig. S5B**). For each neuron population $p$, we used the 2D probability matrix $\mathbf{P}_p(x,y)$ derived from transcriptomics data to sample neuron locations. Specifically, we binomially sampled the 2D spatial density $\mathbf{P}_p(x,y)$ (**Fig. S5C**) to obtain a binary matrix $\mathbf{M}_p(x,y)$ where:

$$\mathbf{M}_p(x,y) \sim \text{Binomial}(1, \mathbf{P}_p(x,y))$$

Here, a value of $\mathbf{M}_p(x,y) = 1$ indicates the presence of a neuron at coordinates $(x,y)$. Each neuron was then assigned a $z$-coordinate by sampling from a uniform distribution over the range $[1,L]$, where $L$ is the network's longitudinal length:

$$(x_i, y_i, z_i) = (x_i, y_i, \mathscr{U}(1,L))$$

For motoneurons, we adapted this approach by using a refined longitudinal distribution $P_z^{\text{moto}}(z)$ based on known motor column locations:

$$z_i \sim P_z^{\text{moto}}(z), \quad \text{for motoneurons}$$

Given a density parameter $D$, we repeated these operations $D$ times in order to control the overall number of neurons in the network. To determine synaptic connectivity, we computed the pairwise distances between neurons in each spatial dimension independently:

$$d_{X,ij} = x_i - x_j, \quad d_{Y,ij} = y_i - y_j, \quad d_{Z,ij} = z_i - z_j$$

We then modeled the probability of a synaptic connection between neurons $i$ and $j$ using Gaussian distributions for each dimension:

$$P_X^p(d_{X,ij}) \sim \mathscr{N}(\mu_X^p, \sigma_X^p), \quad P_Y^p(d_{Y,ij}) \sim \mathscr{N}(\mu_Y^p, \sigma_Y^p), \quad P_Z^p(d_{Z,ij}) \sim \mathscr{N}(\mu_Z^p, \sigma_Z^p)$$

where $\mu_X^p$, $\mu_Y^p$, and $\mu_Z^p$ are the means, and $\sigma_X^p$, $\sigma_Y^p$, and $\sigma_Z^p$ are the standard deviations estimated from axonal projection data. The overall synaptic probability was computed as:

$$\mathbf{P}_{syn}(i,j) = \left(P_X^p(d_{X,ij}) \cdot P_Y^p(d_{Y,ij}) \cdot P_Z^p(d_{Z,ij})\right)^{1/3}$$

This combines the probabilities from each dimension into a single measure of connection likelihood. We then binarized this probability to create a binary adjacency matrix $\mathbf{A}$:

$$\mathbf{A}(i,j) \sim \text{Binomial}(1, \mathbf{P}_{syn}(i,j))$$

where $\mathbf{A}(i,j) = 1$ indicates a synaptic connection between neurons $i$ and $j$. To finalize the connectivity matrix, each neuron population $p$ was assigned a gain parameter $g_p$ (**Fig. S5C**). The final connectivity matrix $\mathbf{W}$ was obtained by scaling the binary adjacency matrix $\mathbf{A}$ with these gains:

$$\mathbf{W}(i,j) = g_p \cdot \mathbf{A}(i,j), \quad \text{for neurons } i \text{ in population } p$$

This approach ensures that each neuron's connectivity is appropriately scaled according to its population-specific gain. The participation of cell types can now be identified as sub population in the projectome, both the ipsi- and contralateral projectome (**Fig. S5D**).

Once the ventral spinal cord network model is constructed, we can carefully examine the network and compare it with empirical observations. First, the cell types are assigned colors and their projection biases are listed (**Fig. S6A**). The locations of the cell somata are distributed in the transverse and longitudinal directions (**Fig. S6B**). The fractional distribution between cell types, both excitatory and inhibitory, as well as their genetic identity are listed (**Fig. S6C**). The connectivity matrix has diagonal lines characteristic for the Mexican hat topology, with local excitation and longer range inhibition (**Fig. S6D**). The network is very sparse (0.97) in agreement with the experimental observations[23]. The normalized density of synapses per cell type is shown in the rostrocaudal direction (**Fig. S6E**). The fraction of cells in space and cell types is also shown (**Fig. S6C**). The axonal trajectories originating from a spot (L6) have a reassuring similarity to the experiment (cf. left and right, **Fig. S6F**). For a fly-through of the experimental tracing see **Video S1**.

### Networks and simulations analyses

#### Principal Component Analysis

Principal component analysis (PCA) was performed on the multidimensional firing rate of the neuronal population. The principal components, denoted as $U_n$ , were obtained as the eigenvectors of the empirical covariance matrix $C$ derived from the $n$ firing rate traces.

$$CU_n = \lambda_n U_n$$

PCA was performed on the $T \times N$ rate matrix using data processing software (`pca()`, MATLAB™). The neuronal population activity is visualized in principal component space over time by projecting the population vector $P$ onto the principal components, resulting in the population vector in the newly defined coordinates $P_{PC}$:

$$P_{PC} = U^T P$$

where $U^T$ is the transpose of the matrix containing the principal components.

#### Oscillation Freqeuncy and amplitude

The oscillatory frequency (measured in oscillations per second, Hz) was estimated by calculating the inter-peak interval within the firing rates projected onto the first principal component.

$$f = \frac{1}{IPI}$$

where $f$ is the oscillatory frequency in Hz and $IPI$ is the average inter-peak interval (in seconds) between successive peaks in the first principal component. Oscillatory amplitude was estimated as the mean root mean square (RMS) over firing rates of all units in a simulation,

$$amplitude \triangleq \frac{1}{N}\sum_{i=1}^{N}\sqrt{\frac{1}{T}\sum_{t=1}^{T} r_{i,t}^2}$$

with $N$ denoting the size of the network, $T$ the simulation duration and $r_{i,t}$ the mean-subtracted firing rate of neuron $i$ at time $t$.

#### Cell type - cell type connectivity

The connectivity between cell types within the network was calculated as the sum of the synaptic weights from all cells of type $i$ to all cells of type $j$, normalized by the total sum of the synaptic weights from all cells of type $i$. This relationship can be mathematically expressed as:

$$C_{ij} = \frac{\sum_{ij} w_{ij}}{\sum_i w_i}$$

where $C_{ij}$ denotes the connectivity metric from all cells of type $i$ to all cells of type $j$, $w_{ij}$ represents the synaptic weight from cells of type $i$ to a specific cell of type $j$, and $w_i$ is the total synaptic weight from all cells of type $i$.

#### Nerve and Muscular readout

The nerve and muscular activity were estimated as the aggregate firing activity of all motoneurons of interest, whether from a pool of motoneurons or from a specific segment.

$$A = \sum_{j=1}^{N} F_j$$

where $A$ represents the total nerve or muscular activity, $F_j$ denotes the firing activity of the $j$-th motoneuron, and $N$ is the total number of motoneurons included in the analysis.

#### Distribution of Premotor interneurons in model

The spatial location of interneurons that project to motoneurons have been an area of interest and experimental investigation[24,25]. The interneurons projecting to the motoneurons were also possible to identify in the model for comparison. We chose two motoneuron pools of samples (tibialis anterior (TA) and gastrocnemius medialis (GM)) since these were also investigated in previous investigations using a retrograde transsynaptic approach (modified Rabies virus)[24,25]. The distributions of TA and GM (**Fig. S7**) appeared realistic and were similar to those seen experimentally[24].

#### Neuronal position and relation to dynamics

One of the proposed principles of rhythm and pattern generation and descending control is the importance of spatial location of cell types in the transverse plane, as well as the shape of the projectome in the longitudinal direction. Hence, an obvious computational experiment to do is to probe the effect of the scrambling position of various neuronal positions.

First, we take the connectivity matrix from above and perform the linear systems analysis procedure to estimate the spatial structure of the emergent dynamics (**Fig. S8A**). Every network instantiation has a connectivity matrix (left). We compute the eigenspectrum of the gain-scaled connectivity matrix (effective connectivity matrix) and identify the dominant eigenvalues (i.e the eigenvalue with the largest real component) which represent dynamical modes of the network (middle panel, **Fig. S8A**). Each eigenvalue has an associated eigenmode (mode) and eigenvector that carries information about the phase and amplitude of every neuron (conceptual sketch of modes 1 and 2 in the third panel). We represent the mode by plotting the eigenvectors in space (right panel). Mode 1 corresponds to the most dominant eigenvalue (i.e. the most amplified dynamical mode of the network). Mode 2 corresponds to the second dominant mode (i.e. the second most likely to be expressed by the network). The first mode expresses alternation between the right and left side. The second mode has right-left synchrony. The distinction can be interpreted as walking (mode 1) and bound locomotion, e.g. gallop (mode 2). The precise right-left phase lag can be adjusted by gain modulation of the contralateral projectome, e.g. by adjusting gain of the commissural projecting cell type V0 (**Video S4** and **Fig. S9C**).

The location of neurons in transverse plane can now be scrambled (cf. **Fig. S8b-C**). Cell type connectivity (middle, **Fig. S8B**), the two dominant eigenmodes (right), and the output of the motoneuron pool (bottom, lateral tibialis and soleus) are shown in the control situation, where they have clear rhythmic alternating activity. When scrambling the position of interneurons in the transverse plane, the cell type connectivity matrix is radically affected (left and middle, **Fig. S8C**). Nevertheless, the first and second eigenmodes still contained alternating rhythms and synchrony (right panel), and the motoneuron pool output looked similar, albeit with reduced amplitude (bottom). Hence, the motor rhythm and pattern generation of the model are remarkably resilient to structural changes in the interneuron population.

However, when the motoneuron pools were scrambled in the rostrocaudal direction, there was a dramatic effect (cf. **Fig. S8D-E**). The location of the motoneuron pools in the lumbo-sacral region of the cord (segments T13-S4) were extracted from anatomical studies[26] and placed in the model (**Fig. S8D**). The spatial pattern of activity in the first mode has a convenient almost one-to-one correspondence with the distribution of motoneuron pools (compare broken and colored lines, right, **Fig. S8D**). The motoneurons receive input like any other spinal neuron, hence from near and far. Since the Mexican hat topology provides mainly excitatory input from local interneurons, the main activation of motoneurons occurs when the bump of activity overlaps with the motoneuron pool. Hence, the large overlap of the bump and the pool results in a clear rhythmic motor nerve activity (bottom). This also means that when the rostrocaudal position of motoneurons is scrambled, the overlap with the propagating bump is poor and the motor nerve output is largely non-rhythmic (**Fig. S8E**). In conclusion, the position of interneurons is not crucial for the generation of rhythms and patterns, whereas the position of motoneuron pools is. Next, the question is what features of the rostrocaudal projections, i.e. the projectome, that is important for rhythm and pattern generation.

### Various forms of projectome and relation to dynamics

The projectome is made up of synaptic projections from various cell types at a given point to various other locations in the spinal cord. We have some general experimental information of the projection directions of various neurons (**Table S1**), but details and statistics are still unknown. Therefore, it is interesting to investigate how sensitive the activity of the model is to various forms of projection patterns - whether there is a unique projection solution to generate the appropriate outcome or whether many different solutions exist. Specifically, we asked whether different projectome features give rise to similar underlying emergent activity? We did this by varying the structure of the projectome in a systematic and objective manner. First, we arbitrarily assign projection bias, that is, having an asymmetric distribution of synaptic targets in the rostrocaudal and mediolateral directions, of a given cell type, and then calculate the dominant activity mode (first and second panel, **Fig. S9A**). Then we randomly changed the projection biases, again calculated the dominant mode and extracted the RC activity profile for left and right hemicord, hence called "hemimodes". This was repeated 500 times, each with a new random projection bias. We preserved the ipsi-contra biases established from the literature. The right side hemimodes were concatenated, plotted, and hierarchically clustered, using UMAP[27] and Louvain method for community detection[28] (third and fourth panel). The hemimodes were then divided into eigtheen clusters, with two highly different clusters selected for illustration (cyan and magenta, **Fig. S9B**). Once the right hemimodes were identified, we performed a subgrouping employing the left hemimodes (**Fig. S9C**). This leads to subclusters with highly similar hemimodes (compare transparent and solid colors, C).

As can be seen (**Fig. S9C right**) projectomes can be radically different, yet generating very similar motor output patterns (cf. Phase plots right). The ipsilateral projectomes are shaped as a Mexican hat (top two projectomes) or have inverted Mexican hat topology (bottom two). The effective dynamical mode in space is visualized with the phase of rhythmic activity for the right and left hemicord (right, shown in gray scale). The difference between the top and the second top is the right-left synchrony (top) and the alternation (second top). As can be seen in the contralateral projectome, the only qualitative difference is that there is a local minimum in the contralateral projectome. For the magenta case, the shape is an inverted Mexican hat in the ipsilateral projectome, and the shape of the contralateral projectome have different options: If the contralateral projectome has a local maximum, there is right/left alternation, whereas if there is a local miminum there is right/left synchrony. Hence, we conclude that rhythm and patterning can be achieved in two ways: First, rhythms can be generated by a Mexican hat or an inverted Mexican hat projectome. The right / left synchrony can be achieved if the local extremum at the origin of the contralateral projectome is pointed in the same direction as the ipsilateral projectome, regardless of whether it is a Mexican hat or an inverted Mexican hat. Similarly, a right/left alternation arises if the local extremum of the contralateral projectome points in the opposite direction to the ipsilateral projectome.

Next, we investigate how the topology of the Mexican hat can be realized. We tried different combinations of cell types in different network models. It turned out that there are multiple solutions to form an ipsilateral projectome shaped like a Mexican hat, with a local maximum in the controlateral projectome (**Fig. S9D**). Both combinations are a distinctly different combination of cell types yet the generated rhythm and pattern are qualitatively similar (cf. right-phase hemimodes). This illustrates an important point: The rhythm and pattern generation is robust to various configurations and not dependent on cell types per se, but crucially dependent on the spatial projection profile of the whole population. What is the role of cell types then? As we will see below, it is about descending control (see below Section "Roles of descending control of motor activity").

Finally, we try a non-Mexican hat shape, the symmetric purely excitatory ipsilateral projectome with the symmetric reciprocal inhibitory projectome (**Fig. S9E right**). Such a projectome was incapable of generating rhythms or patterns in the rostro-caudal direction. This suggests that two features are important for rhythm- and pattern generation: First, there must be an asymmetry in the rostrocaudal direction. Second, there must be either a Mexican hat topology (although with asymmetry) or an inverted Mexican hat topology. Looking closer at the latter case, while an inverted Mexican hat is able to generate rhythms and patterns, it is difficult to slow down and stop the bump of activity, because sustained local activity is unstable. Therefore, we suggest that vertebrate species that have to be able to arrest the progression of movement, for instance, a cat stopping the locomotion when seeing a mouse, is only possible if the projectome has a regular Mexican hat topology. In other invertebrates species where motor arrest is not required for any posture, for instance a swimming movement where the stop has a straight spinal cord, an inverted Mexican hat topology could also generate appropriate motor behaviors. However, since the regular Mexican hat topology can serve movements in any species, we propose this as the most universally conserved.

In conclusion, this analysis first shows that the asymmetrical Mexican Hat topology is the most common profile to produce rhythms and patterns among random generated networks. Second, while the longitudinal position of motoneurons is crucial for proper motor output, the position of the interneurons is not crucial. This raises an intriguing question: is the position of interneurons important for something else and, if so, what? As we shall see, the position of the internurons is important for the descending control of all aspects of motor generation.

### Role of interneuron position and descending control of motor generation

To investigate the role of the position of interneurons in the transverse plane we consider two situations: first, the situation where the position is based on empirical data, the single cell RNA sequencing and spatial transcriptomics (control condition, (**Fig. S10A**)); second, a condition in which the soma positions are randomly scrambled (**Fig. S10B**). The cell type correlation matrix in the two situations has the same configuration as earlier (middle, **Fig. S8B**). The change in cell type connectivity is visible by having more even distribution, except for the connection to motoneurons, which is kept (vertical black row). Next, we provide two types of descending input: A well-known descending excitatory input (associated with vGlut2), which comes from the caudal brain stem[29] (the lateral paragigantocellular nucleus, LPGI, middle panel). When the positions of cell types were scrambled, rhythm and pattern generation was disrupted and replaced by a tonic activity (cf. A and B, middle **Fig. S10**). The cell type target specificity is indicated by a half-circle polar distribution. When the descending input was aiming globally, the scrambling had no effect (cf. bottom, **Fig. S10A-B**). This observation delineates a possible role for spinal cell types. The descending control of motor activity is only possible due to the spatial segregation that cell types represent combined with their specific and dissimilar rostrocaudal projection patterns.

### Modifying projectome by descending input and hence gait

The projectome is defined as the sum of the excitatory and synpatic input as a function of the rostrocaudal distance from a given point. If the gain of a certain cell type is changed by e.g. descending input, the firing rate of this cell type may change and hence the synaptic input to the target (**Fig. S11A**). This can be viewed as a change in the connectivity, also called "effective connectivity". Once the connectivity is changed to a new effective connectivity by gain modulation of individual cell types by descending input, the projectome is also changed and therefore also the eigenvalue spectrum and hemimodes (right, **Fig. S11A**). To investigate this further, we considered three different values of gain (**Fig. S11B**). Gain modulation in this model is accomplished by descending input from glutamatergic neuron in the caudal brain stem (LPGi) or serotonergic cells located in the caudal Raphe nucleus (bottom). The population dynamics is simulated with a ramping up of these descending fibers (**Fig. S11C**). The size of the effective projectomes increases as the gain increases (second-bottom panels), which also results in a faster frequency and wave propagation due to stronger asymmetry in the projectomes. The eigenvalue spectra for the three situations also demonstrate that there are two dominant modes, which swab places as the descending drive increases (arrows). The consequence is that the right-left activity switches from alternation to synchrony (cf. hemimodes, bottom-right).

### Stability of the rhythmic activity

An important issue with modeling is how much the parameters of the model need to be optimized to induce a certain behavior. In our case, behaviors are the generation of rhythms and patterns. The issue to investigate is the sensitivity to the length of axonal projections and the spread of the projections (left and middle panels, **Fig. S12A**). Next, we define two metrics to characterize the effect of the parameters and hence the robustness of the model (**Fig. S12A**). First, we define the distance of the eigenvalue from the linear stability line as the "Margin of stability", **M**. Second, we define the variability of the position of the dominant mode normalized by the position, $\Delta R/R$. Using these two metrics, we first inspect the effects of neuronal density, i.e. how densely populated the cord is, on the robustness of the modes. As the density increase, the variability decrease and the margin of stability increase (**Fig. S12B**). Three sample densities are shown (right). Hence, having more cells in the cord generally stabilizes dynamical modes, which perhaps could explain the large number of spinal interneurons that participate in the relatively simple low-dimensional rhythmic locomotor activity. Finally, we inspect the impact of the width of projection distributions on the robustness. There is an optimal in both the variability and the margin of stability. The variability has a minimum around 700$\mu$m (red bar), which is the same place where the margin of stability has a maximum (**Fig. S12C**). This optimum depends on the density. Spinal network instantiations shown for three projection widths (100, 900, and 1700 $\mu$m, right).

### Mathematical Note: Spectral Analysis

Spectral analysis of the connectivity matrix **W** allows prediction of network dynamics. To see how, consider our rate-based network model

$$\dot{\mathbf{x}} = -\mathbf{x} + \mathbf{W}F(\mathbf{x}) + \mathbf{I_{ext}} \tag{1}$$

where $\mathbf{x}$ is a vector that holds the momentary inputs of the $N$ neurons, $\dot{\mathbf{x}}$ its time derivative, $F$ the neuron gain function mapping the input to the firing rate and $\mathbf{I_{ext}}$ is the vector of external inputs. For simplicity, the variables are unitless and the system time constant is set to 1. For balanced networks with constant, homogeneous external inputs, a uniform fixed point solution to (1) exists where $\mathbf{x}^* = \mathbf{I_{ext}}$.

The dynamics of a small perturbation $\delta(t) = \mathbf{x}(t) - \mathbf{x}^*$ follows the dynamics:

$$\dot{\delta} = -\delta(t) + \mathbf{W}F(\mathbf{x}^* + \delta(t)). \tag{2}$$

Taylor expanding around $\mathbf{x}^*$ yields

$$\dot{\delta} = -\delta(t) + \mathbf{W}(F(\mathbf{x}^*) + F'(\mathbf{x}^*)\delta(t)) \tag{3}$$

where we've neglected higher order terms. Since $\mathbf{W}F(\mathbf{x}^*) = 0$ , we obtain

$$\dot{\delta} = -\delta(t) + g\mathbf{W}\delta(t) \tag{4}$$

where we identify $g$ as the gain, i.e. the slope of $F$ evaluated at the fixed point $\mathbf{x}^*$. Rearranging the now linear (4), we arrive at the standard form

$$\dot{\delta} = \mathbf{A}\delta \tag{5}$$

where $\mathbf{A} = g\mathbf{W} - \mathbf{I}$ and $\mathbf{I}$ is the identity matrix. Previously, $\mathbf{A}$ was referred to as "the effective connectivity matrix"[8], since it is the connectivity matrix scaled by the neuronal gain. As with any system of linear ordinary differential equations, a general solution to (5) is given by

$$\delta(t) = c_1 e^{\lambda_1 t}\mathbf{v}_1 +, ..., + c_N e^{\lambda_N t}\mathbf{v}_N \tag{6}$$

where $\mathbf{v}_i$ is the $i_{th}$ eigenvector of $\mathbf{A}$ and $\lambda_i$ its corresponding eigenvalue. By examining the spectrum of eigenvalues belonging to the connectivity matrix $\mathbf{W}$, we can predict the response of the network to perturbations in the uniform state. If $\Re(\lambda_i) < 1$ for all $i$, all perturbations will decay in time and the fixed point is stable. If the leading mode has $\Re(\lambda_{lead}) > 1$, the fixed point is unstable and any perturbation along $\mathbf{v}_{lead}$ will grow in time and characterize the dynamics. Finally, if the fixed point is unstable and $\Im(\lambda_{lead}) \neq 0$, the perturbation response will oscillate, leading to rotations in neural space.

**Table S1** Experimental information on cell types (most left colomn), rostrocaudal, dorsoventral and mediolateral projections. The approximate location of the somata in the spinal layers (right) and their references (most of the right column).

| Table S1: Cell type Projections and References | | | | | |
|---|---|---|---|---|---|
| Population | RC | DV | ML | Layers | Ref. |
| Chx10/V2a-1 | Local | Ventral | n/a | VI, VII, VIII & IX | 1,30–32 |
| Chx10/V2a-2 | bidirectional, caudal bias, 5000 $\mu m$ | Ventral | n/a | VI, VII, VIII & IX | 1,30–32 |
| V1 | Rostral bias | Ventral | n/a | VI, VII, VIII & IX | 33,34 |
| V2b | Caudal bias | Ventral | n/a | VI & X | 32 |
| V3 | Caudal bias, commissural | n/a | n/a | VII, VIII & IX | 35,36 |
| DI6 | Caudal bias, 3 segments, 4000 $\mu$m | Ventral | n/a | VI, VII, VIII & IX | 37–39 |
| V0d | Local, commissural | Ventral | n/a | VIII | 37 |
| V0v | Rostral bias, 1-4 segments, commissural | Ventral | n/a | VIII | 37,40 |

**Video S1** **Interneuron projections in the mouse spinal cord** Tissue cleared (iDisco+) mouse spinal cord imaged on a light sheet microscope (Ultramicroscope Blaze, Miltenyi Biotec). A virus had been injected in the lumbar spinal cord at one spot to nonspecifically and sparsely infect the interneurons there, for the purpose of partially tracing the projectome. The cells that expressed fluorescent reporter, have their fluorescence enhanced with immunohisochemistry after clearing. Many axons appear bright fluorescent. Many cross over and both ascend and descend. Other fibers stay ipsilaterally and also ascend or descend. Since the labeling is sparse, there are many more projection fibers than are visible in the video.

**Video S2** **Spinal network simulation demonstrating the slowdown and halt of rhythmic activity.** Here a one-dimensional network model and simulation of the activity are shown. The projectome is composed of five subpopulations, each representing one of the five lobes of the projectome. The top left panel shows the firing rate activity of all neurons at a given longitudinal location. The top right panel shows the projectome, i.e. the net synaptic input to locations from fibers originating from a given location, i.e. at the origin. The lower five panels represent the firing rate activity of subpopulations 1-5 in the shaded region in the top left panel. The open space in the histograms is due to other cell types taking these locations. First, there is a global input of constant input, resulting in rhythm and pattern generation. Some time later, the gains of some the subpopulations are modulated: The subpopulation 2 (corresponding to V1) is decreased and the subpopulation 4 (corresponding to V2b) is increased. As a consequence, there is a slow-down of the rhythm spatial propagation.

**Video S3** **Building the projectome out of cell types** The rostro-caudal projection of different cell types constitutes the projectome. Here are some of the cell types that make up the spinal projectome. First, DI6, then motoneurons (MN), then V0d, V0v, V1, V2a-1 (short-range projecting) and V2a-2 (long-range projecting), and V3. In the end, the projection pattern of all of them is shown in the projectome.

**Video S4** **Network model of rodent spinal cord and simulation of motor activity** The right and left side of the lumbar spinal cord of a mouse is modeled and simulated. First, all cells receive a global constant input, which is transformed into spatiotemporal rhythms and patterns. The dots represent the neuronal soma and the size of the dots represents the firing rate of that neuron. Bumps of activity emerge at the rostral end and travel toward the caudal end. This occurs in both the right and left side of the spinal cord, although in alternation due to and orchestrated by the commissural connections. Half way through the simulation, the gain of a subset of commissurally projecting interneurons, V0, is changed. As a consequence, the right-left rhythms are now synchronized. The right-left phase difference can assume any value depending on how much the gain is modulated. Here, a phase difference of zero is shown at the end.

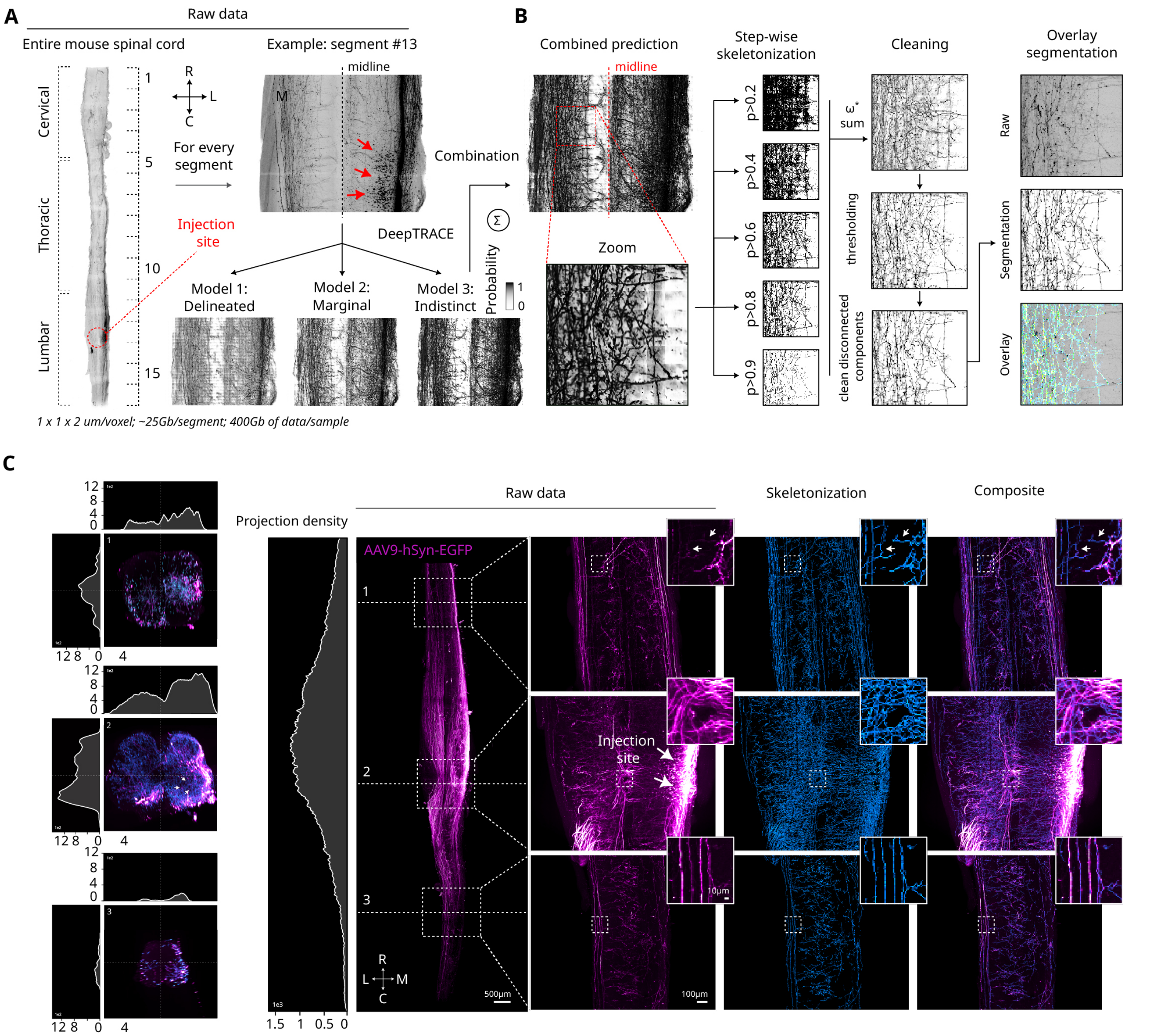


**Fig. S1.** | **Mapping of projections from a specific location using virus injection in the mouse spinal cord. A**, On the left, a top view of the raw data from an entire spinal cord stained for red fluorescent protein (RFP) is shown, with the injection site highlighted in red (red arrows point to RFP+ cells). Due to the large data size ( 400GB), the segmentation pipeline was run on segments of the data ( 25GB per segment). Each model in the pipeline is sensitive to different aspects of the data: Model 1 is more sensitive to well-delineated structures, Model 2 to marginal objects, and Model 3 to indistinct and diffuse signals. **B**, software (DeepTRACE) predictions from the models are then combined, as previously described[1]. A stepwise skeletonization is applied to the combined predictions. The binary skeleton outputs are weighted based on the initial probability thresholds used in their generation and subsequently summed. This approach preserves prediction confidence while avoiding the fragmentation of connected structures due to threshold-based segmentation. Truncated and disconnected objects are then removed to produce the final segmentation mask. The resulting segmentation mask aligns well with the raw data, although some projections remain undetected (indicated by red arrows). **C**, The "skeletonization" panel demonstrates the segmentation quality along the entire spinal cord for an example sample (entire spinal cord stained for GFP is shown). The projection density graphs show the segmented voxel count along the rostro-caudal, dorso-ventral, and medial-lateral axis (proxi of the projection density). While most projections are localized near the injection site, many are detected at the extreme ends of the spinal cord. Additionally, a noticeable ipsilateral/contralateral bias in projection distribution relative to the injection site is observed, though further analysis is required to fully characterize these differences.

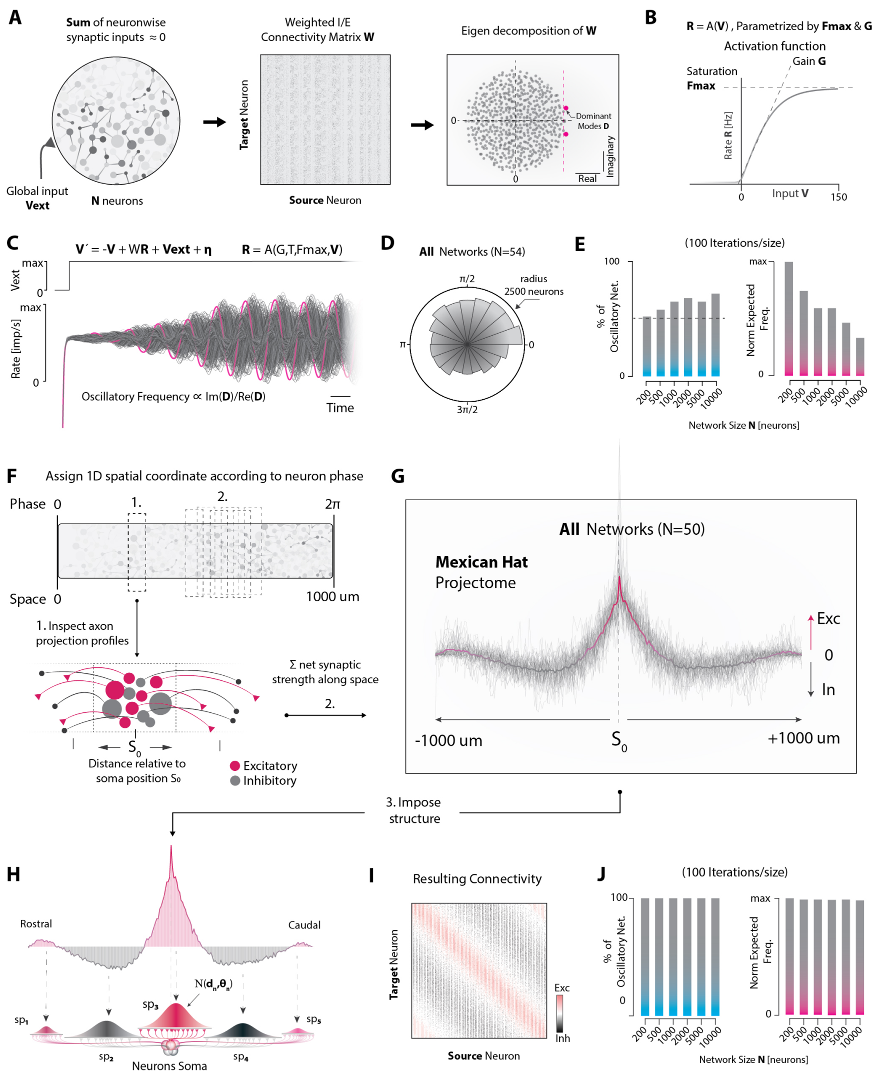


**Fig. S2.** | **Spatial embedding of balanced sequence generator networks reveals a Mexican Hat projectome. A**, Example of a balanced sequence generator (BSG), consisting of N randomly coupled neurons with balanced excitatory and inhibitory recurrent inputs receiving a global external input. The network $N \times N$ connectivity matrix **W** has a circular spectrum of eigenvalues, and a dominant complex mode (red) beyond the stability line (red dashed), predicting sequence-generating properties. **B**, The synaptic input to each neuron is converted to a momentary firing rate by a hyperbolic tangent activation function. **C**, When subjected to a global drive and simulated according to dynamical equations in the firing rate formalism, BSG networks oscillate in time with a frequency proportional to the imaginary part of the dominant eigenvalue. **D**, BSG networks oscillate with a quasi-continuous distribution of phases, a phenomenon termed "rotational dynamics". **E**, Of random, balanced networks, only approx. half exhibit oscillatory properties across sizes N = [200, 500, 1000, 2000, 5000] (100 networks/size) (left). Predicted frequency tends to decline as BSG networks grow (right). **F,G**, BSG networks (n=50) were embedded on a linear space according to a phase-space coherence rule, such that the phase-position of a neuron in an arbitrarily chosen reference period reflected its position on the line. From the connectivity and spatial positions, the net synaptic strength over the relative distances revealed an asymmetric Mexican Hat projectome with local excitation and broader inhibition. **H,I**, The unfolded projectome was recapitulated in spatial 1D networks through five subpopulations, each with a projection bias matching one of the five lobes of the Mexican Hat. This yielded a spatially ordered connectivity matrix where excitatory (red) and inhibitory (black) synapses were visible as diagonal bands. **J**, Across sizes, all networks (100 networks/size) were predicted to oscillate with stable frequency.

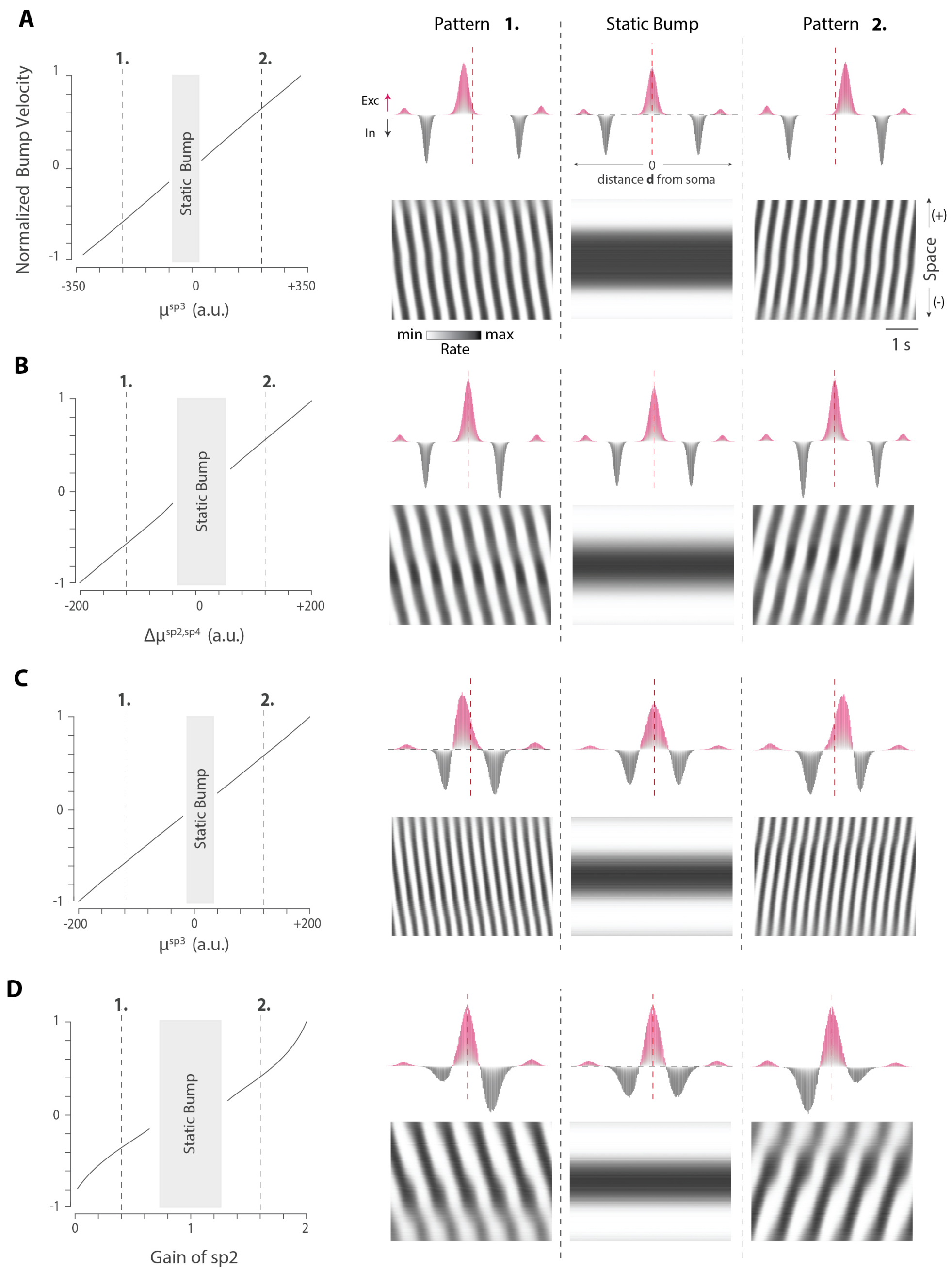


**Fig. S3.** | **Different projectome asymmetries efficiently controls direction and velocity of wave propagation in 1D networks A**, 1D networks (n=34, size=2000) constructed according to **Fig. S2H** with non-overlapping projection distributions for the five subpopulations. As the mean projection distance of the local excitatory subpopulation, $\mu^{sp3}$ , was moved away from zero point, the velocity of wave propagation (oscillatory frequency) linearly increased with a propagation direction reflecting the bias direction (left). When $\mu^{sp3}$ approached zero, yielding a near symmetric projectome, wave propagation terminated and a static bump pattern characterized the dynamics (right). **B,** Networks (n=21, size=2000) constructed as in **A**. As $\mu^{sp2}$ and $\mu^{sp4}$ (left and right projecting inhibition, respectively) were shifted together away from symmetry, propagation velocity increased with a direction that reflected the direction of least inhibition (left). Again, at symmetry, only the static bump emerged (right). **C,** Networks (n=21, size=1000) with overlapping projection distributions for subpopulation 2, 3 and 4. Bump velocity as a function of $\mu^{sp3}$ and the spatiotemporal pattern matched that in **A** . **D,** Networks (n=100, size=1000) constructed with overlapping projection distributions, as in **C**. Instead of varying projection biases, asymmetry was induced by modulating the weight of sp2 synapses (and sp4 synapses, inversely). Bump velocity was controlled near linearly by gain modulation with propagation direction reflecting the direction of least inhibition.

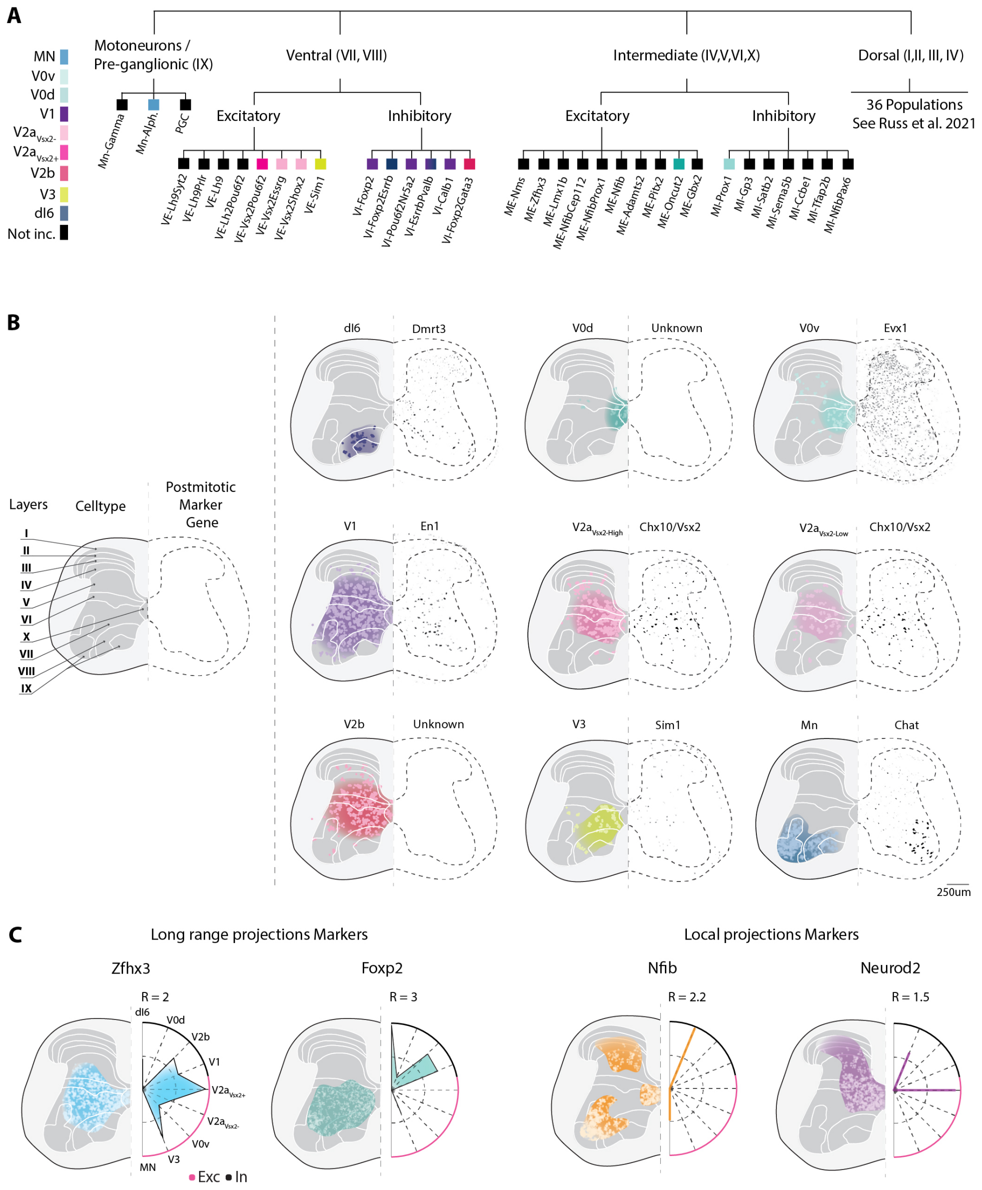


**Fig. S4.** | **Spatial transcriptomics delineates the transverse location of cardinal cell types A**, Tree representation of all ventral cell types from an harmonized cell atlas.[20] . The coarse cell types considered in this study are highlighted in colors and correspond to group of subtypes from similar developmental lineage. **B**, Lamniar organisation of cell types in the mouse lumbar (L4) segments. For each spinal cord representation, the left side corresponds to the transverse location of the specified cell type, derived from deconvolution of spatial transcriptomics with single cell RNA sequencing (see Methods). The right side shows the location of the primary marker gene of the cell type derived from in-situ hybridization (Allen Mouse Spinal Cord Atlas). **C**, Differential expression of marker genes for long range and locally projecting neurons[22]. Each panels shows spatial regions where genes are differentially expressed, on the left, and a polar plot of relative gene expression level for each cell type, on the right. Radius size is indicated for each plot. Central dashed line is half-radius. The spatial regions are in close agreement with those reported previously[22]. This analysis suggests specific projection patterns for most of the cell types considered. DI6, V2b, V1, V2a , V0v and V3 express long ranges marker. V2a-Vsx2+, MN and V0d express short ranges markers. This is in good agreement with exsiting morphological data (see Table S1) and corroborates most projections used in our model, except for V3.

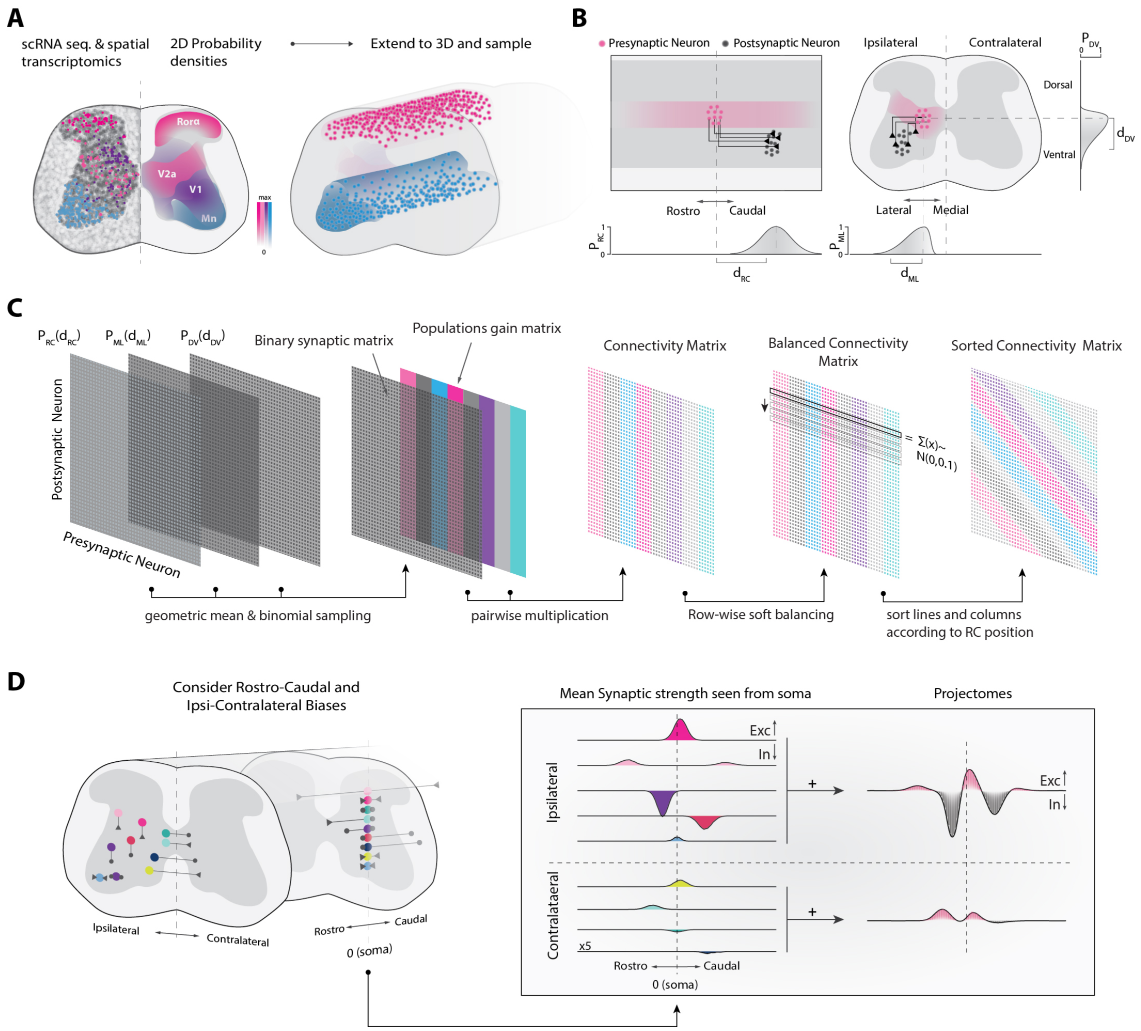


**Fig. S5.** | **Construction of ventral spinal cord motor network connectivity and projectome from empirically inferred cell type location and projection biases.** **A**, The cell type spatial probability distributions in transverse plane are derived from combined single cell RNA sequencing and spatial transcriptomics in mouse lumbar spinal cord (L5). Somas of cell types are generated from these probability distributions and extended in the longitudinal axis. **B**, neurons of a given cell type (presynaptic neuron, red) are targeting other cells of various identity (postsynaptic neurons, black) based on their rostrocaudal (RC), mediolateral (ML) and dorsoventral projection probability, which means ($d_{RC}$, $d_{ML}$, and $d_{DV}$) are inferred and estimated from literature. **C**, The effective connectivity matrix is formed first by binomial sampling of the spatial probability matrices which is then multiplied by the population gain matrix, where different cell types can be modulated by descending input (colors). Next, the weights are adjusted to maintain a row-wise balanced of excitation and inhibition. The matrix row and columns are sorted to match the longitudinal location (right). **D**, Different cell types are located at their preferred transversal locations, and they have their preferred projection directions. The longitudinal projections can be collapsed and divided into ipsi- and contralateral projections from a given spot (vertical broken lines, right), depending on their cell type (colored distributions). The sum of excitation and inhibition for a given set of gains constitutes the effective ipsi- and contralateral projectomes (right).

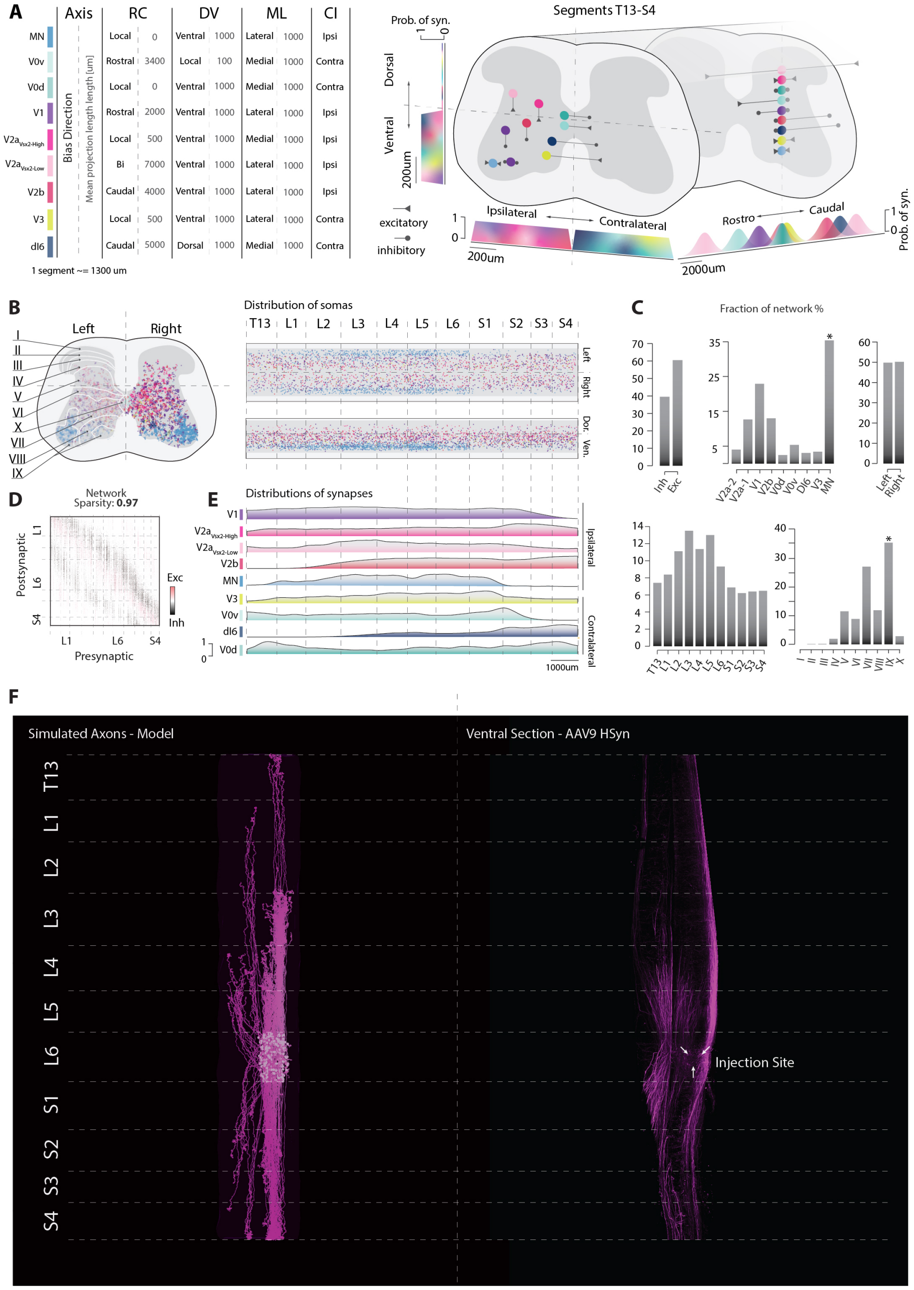


**Fig. S6.** | **Spatial features of the spinal model A**, Left: Table of projection biases on the 3 axis of space (Rostro-Caudal RC, Dorso-Ventral DV, medio-lateral ML and Contr-Ispi lateral CI) for each population. Right: Spatial representation of projection biases and terminal likelihood distributions. **B**, Laminar and longitudinal neuron distributions in the model. **C**, Fractions of neurons transmitter types (Excitatory/Inhibitory), celltypes, and spatial locations (left/right, segments and laminae). A star indicates where values are intentionally overestimated for technical reasons requiring inclusion of a larger number of motoneurons in the model. **D**, Connectivity matrix and network sparsity. **E**, Normalized densities of synapses per celltype along the rostro-caudal axis. Synaptic densities are non-uniform as a consequence of the projection biases. **F**, Left: Simulated axonal projections emanating from a single segment (L6) in the model. Right: Cleared spinal cord showing viral axonal tracing of all neurons infected at the injection site (AAV9 HSyn promoter).

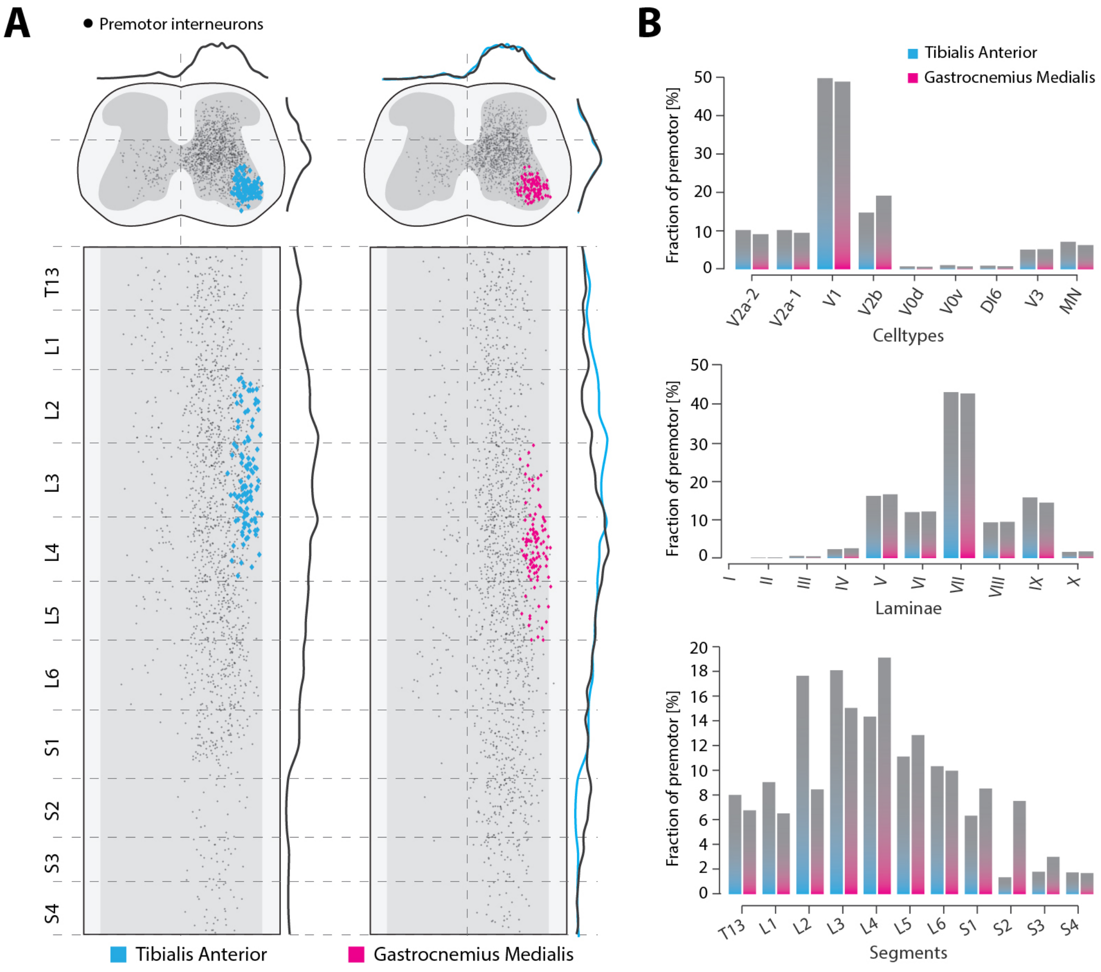


**Fig. S7.** | **Motoneurons and premotor neurons in the model. A**, Two sample motoneuron pools, Tibialis Anterior (left, blue) and Gastrocnemius Medialis (right, pink), in transverse plane (top) and longitudinal plane (bottom). Premotor interneurons are indicated as black dots. Their spatial distributions are shown above and to the right side of each panel. The blue lines in the right panel are copies of the premotor distributions of Tibialis Anterior, for direct comparisons. **B**, Fractions of celltypes, laminar and segmental distributions composing the premotor populations of each motor pool.

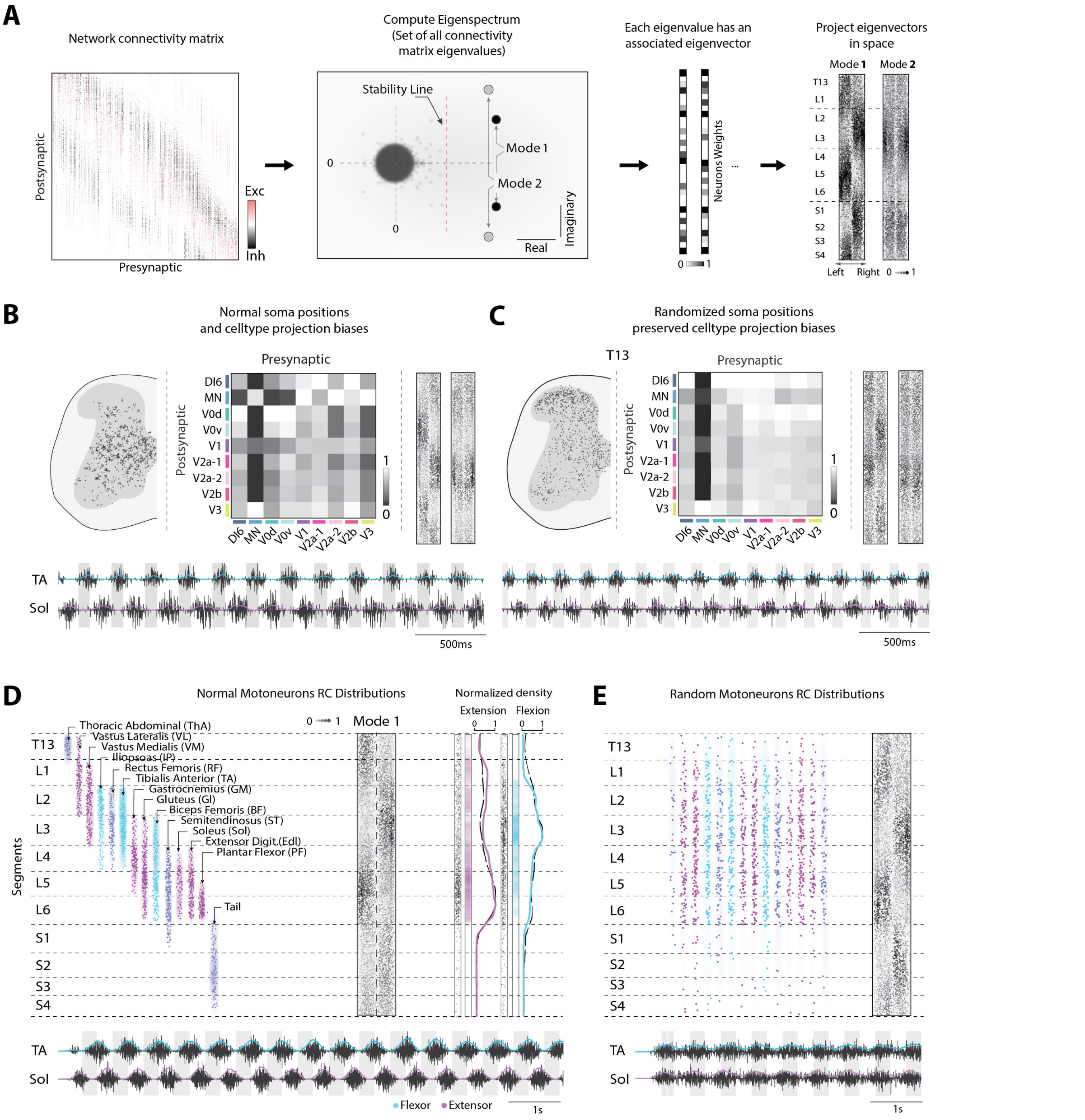


**Fig. S8.** | **Impact of neurons positions on emergent dynamics A**, Procedure to estimate the emergent dynamics and spatial activity: From the connectivity matrix (left) the eigenspectrum is computed (2nd panel) to identify the dominant eigenvalues (i.e with the largest real value). The associated eigenvector with values of every neuron (eigenvector 1 and 2 sketched in third panel). These eigenvectors are mapped onto the spinal network to show spatial activity (Mode 1 and 2). Mode 2, i.e. the second most likely to be expressed by the network, has right-left (right panel). **B**, For networks with projection biases and length as established in Fig. 3 and Fig. S6, the linear analysis for normal (i.e. from spatial transcriptomics) interneurons (i.e. non-motoneurons) somas distributions predicts a dominant left-right and flexion extension alternation. A second instantiation, where somas position are randomized in the whole grey matter is shown in **C**. Cell type specific spatial projection biases are preserved but the resulting cell type to cell type connectivity is altered. Nevertheless, the predicted dominant mode preserve their structure along the rostrocaudal axis. **D**, The linear analysis for normal motoneuron pools distributions (left side) predicts left-right and flexion-extension alternation **E**, For a randomized position of the motoneurons, the predicted dominant mode remains unaltered but the muscular output show poor rhythmicity and alternation.

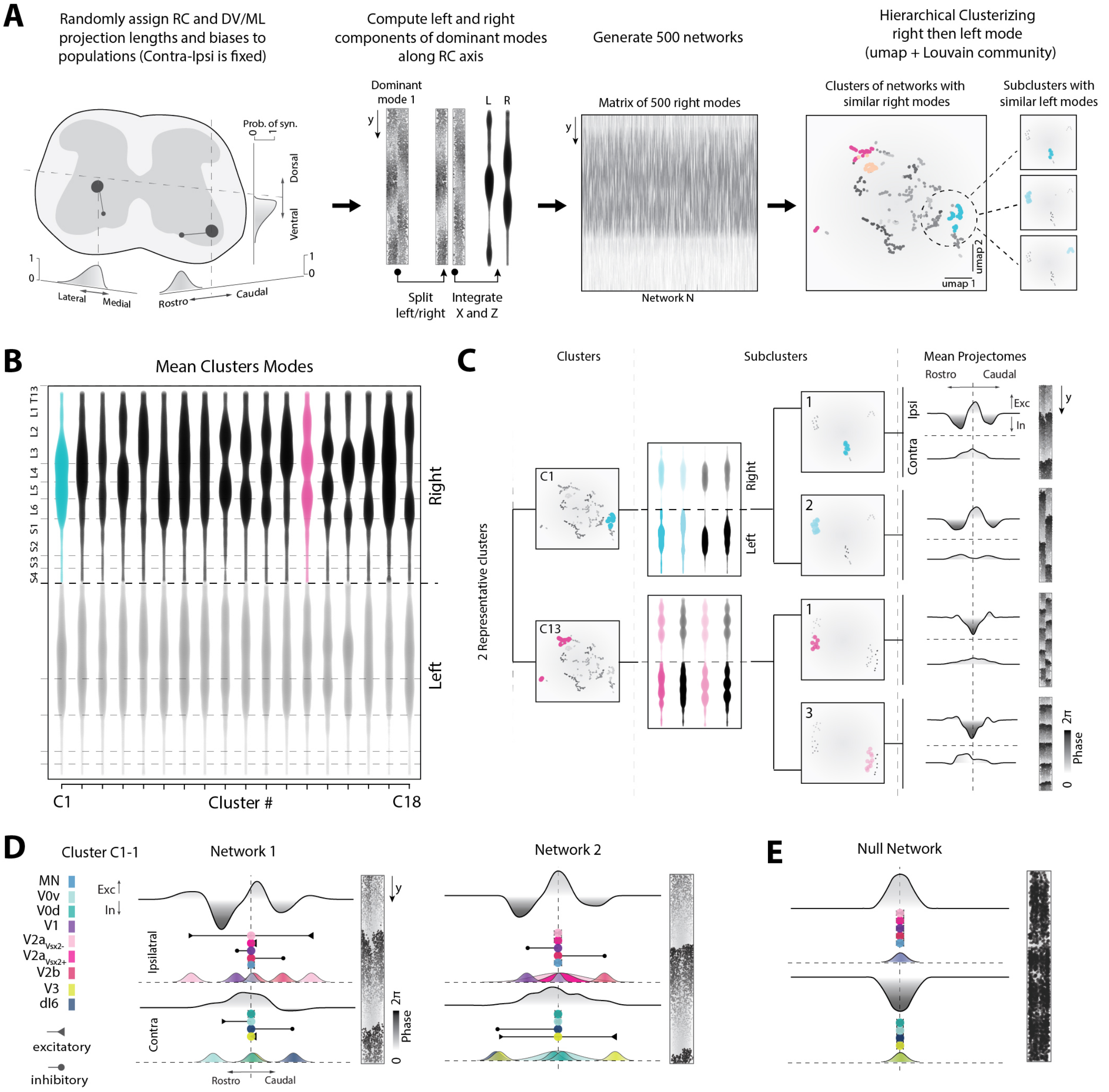


**Fig. S9.** | **Impact of projectome structure on emergent dynamics. A**, Unbiased procedure to identify projectome features underlying similar emergent dynamics. We randomly generate 500 networks in which we arbitrarily assign rostro-caudal and medio-lateral biases (directions) and lengths (means of synaptic distributions, in a [0, 5000um] range) to celltypes . We preserved the ipsi-contra biases established from literature. For each network, we predict the spatial structure of the dominant mode following the procedure in Fig S8A. We separate the left/right components and collapse the hemimodes along the rostro-caudal axis (L and R). We concatenate all rights hemimodes in a matrix. We reduce the dimensionality of this matrix to 2 dimensions using UMAP. In the 2D space, we identify clusters of network with similar right hemimodes and proceed to a subclustering employing the left hemimodes. This leads to subclusters with highly similar dynamical modes in space. **B**, Diversity in spatial structure of collapsed dynamical modes after clustering on the right hemimodes. Two highly different clusters are highlighted (blue and pink). **C**, Left: UMAP representation of the 2 selected clusters of panel B (C1 and C13). Middle: Spatial structure and UMAP of the subclusters of C1 and C13. Selected subclusters are highlighted. Right: Mean projectomes (ipsilatareral and contraleteral components) and effective dynamical mode in space. **D**, Sample of 2 networks from subcluster C1-1 with similar projectome shape and dynamical mode, but dissimilar celltype projections. **E**, Example of a network without a projectome (i.e. null network, all celltypes project locally). Linear analysis predicts no emergent structured dynamics and all cells are synchronized.

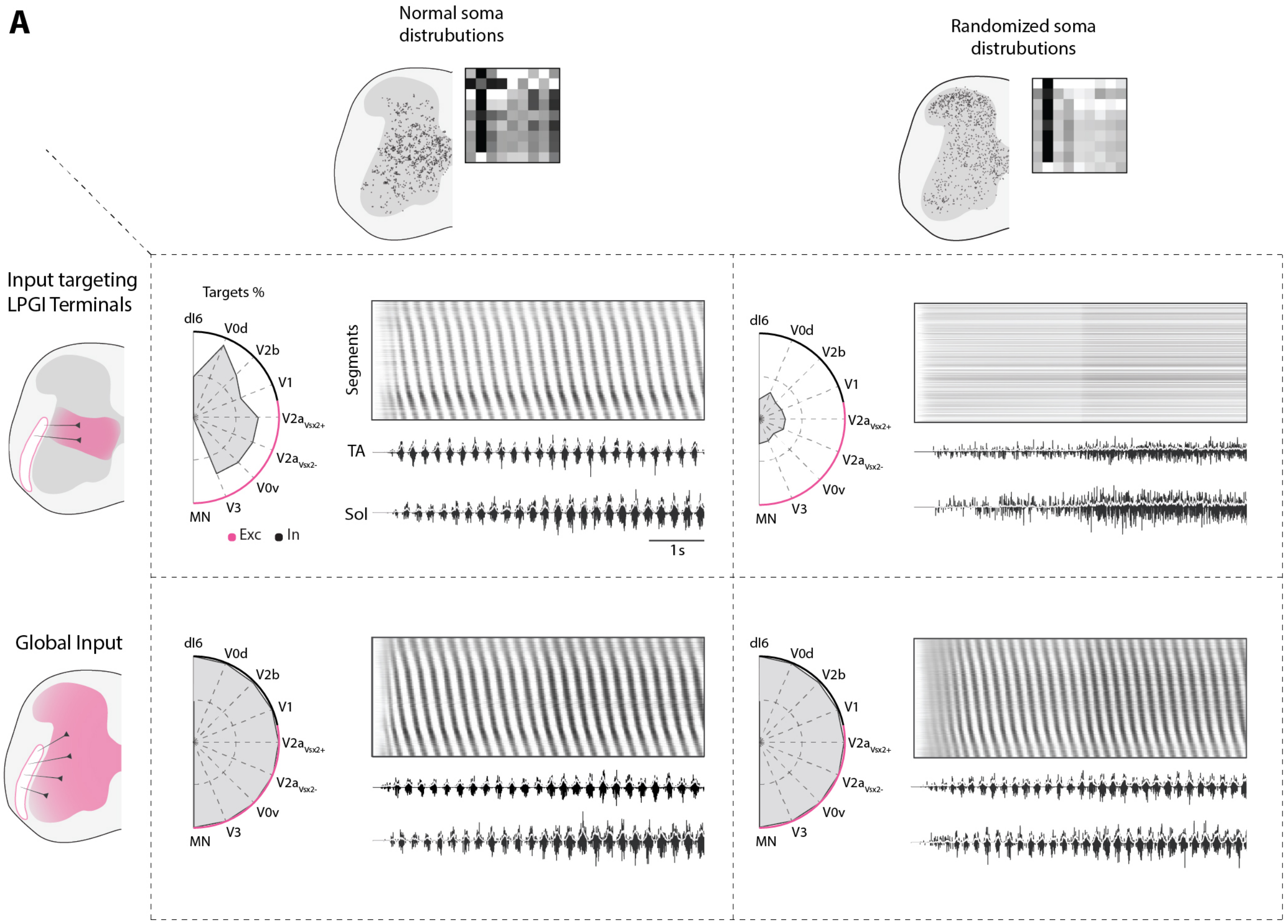


**Fig. S10.** | **Impact of cell type position in the transversal plane on descending control. A**, Columns illustrate the transversal cell distribution and its corresponding population-based connectivity matrix for the nine populations modelled in the case of normal (left) and randomized (right) cell positions. Rows illustrate the areal distribution of descending lateral paragigantocellularis (LPGI) terminals (top) and global input terminals (bottom). When the transversal cell distribution is normal, LPGI fibers synapse onto a large fraction of excitatory and inhibitory classes and external drive from these fibers is sufficient to initiate spatiotemporal sequences down the rostrocaudal spinal axis as well as flexor-extensor alternation in soleus (Sol) and tibialis anterior (TA) muscles. When neuron positions are randomized, only a small fraction of cell populations receive input from LPGI fibers, and descending drive is insufficient to generate sequences or locomotor-like muscle activity. When the descending input is global however, all neurons receive external drive and sequences emerge for both normal and randomized transversal positions (bottom row).

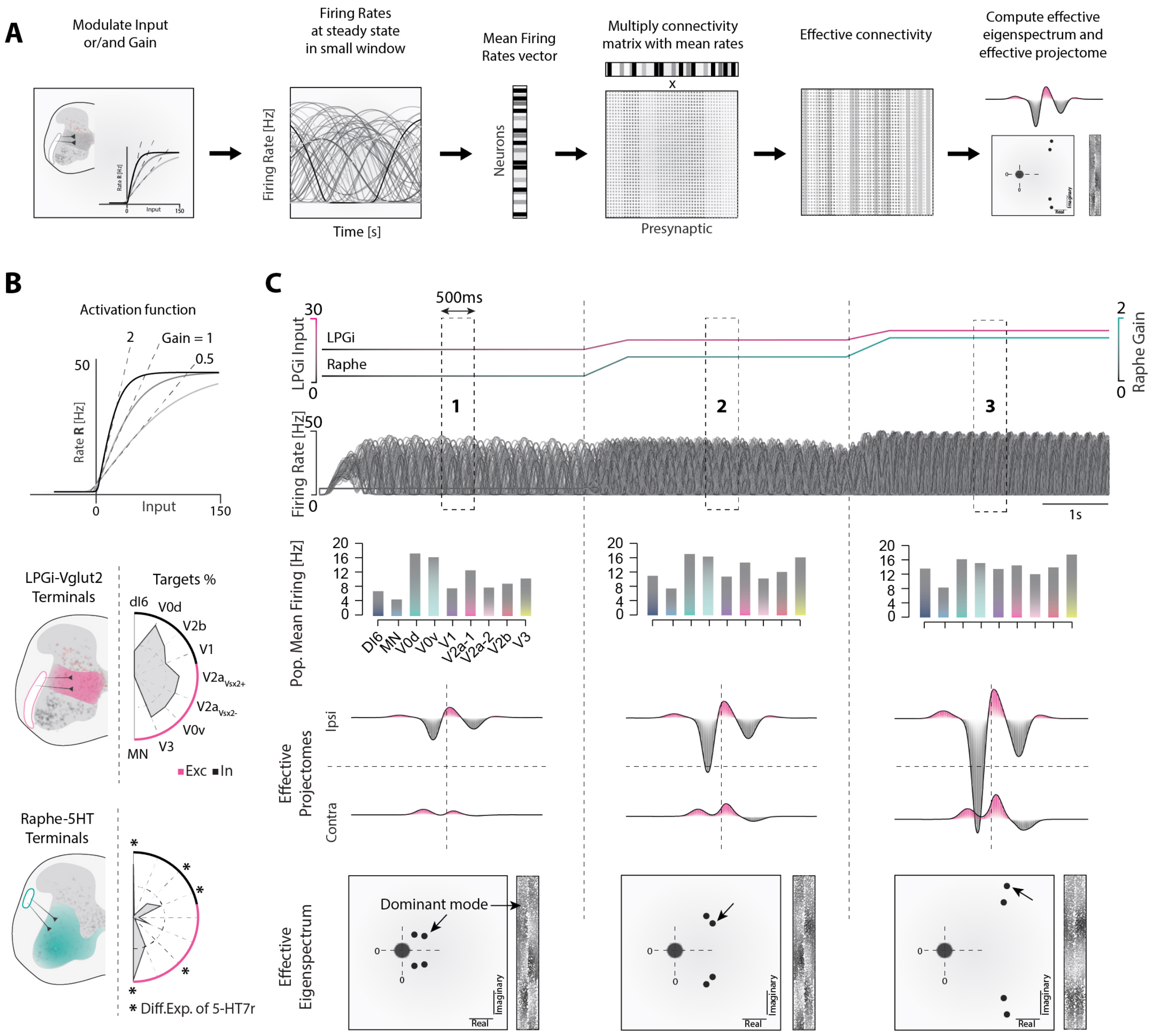


**Fig. S11.** | **Impact of input and gain modulation on the projectome and emergent dynamics. A**, Procedure to continuously estimate an effective projectome when varying input parameters. In simulations, we modulate the gain and/or the input to the network. This effectively results in changes in firing rates. We apply a centered sliding mean of 500 ms to each neurons to estimate the mean firing rate at all times. For one window, this results in a vector of length N (i.e. the number of neurons in the network). We multiply the columns of the connectivity matrix with the mean rate vector, elementwise. The result is an effective connectivity matrix at a time t. We compute the spectrum and the projectome of this matrix. **B**, Top: the activation function (hyperbolic tangent, see Methods) of the neurons in the model. Middle: Location of the terminals of the LPGi fibers and targeted cell types. Bottom: Location of the terminals of the Raphe Obscurus fibers and targeted cell types. Radius is 100 percents. Stars indicate a significant differential expression of 5-HT7 receptor. **C**,

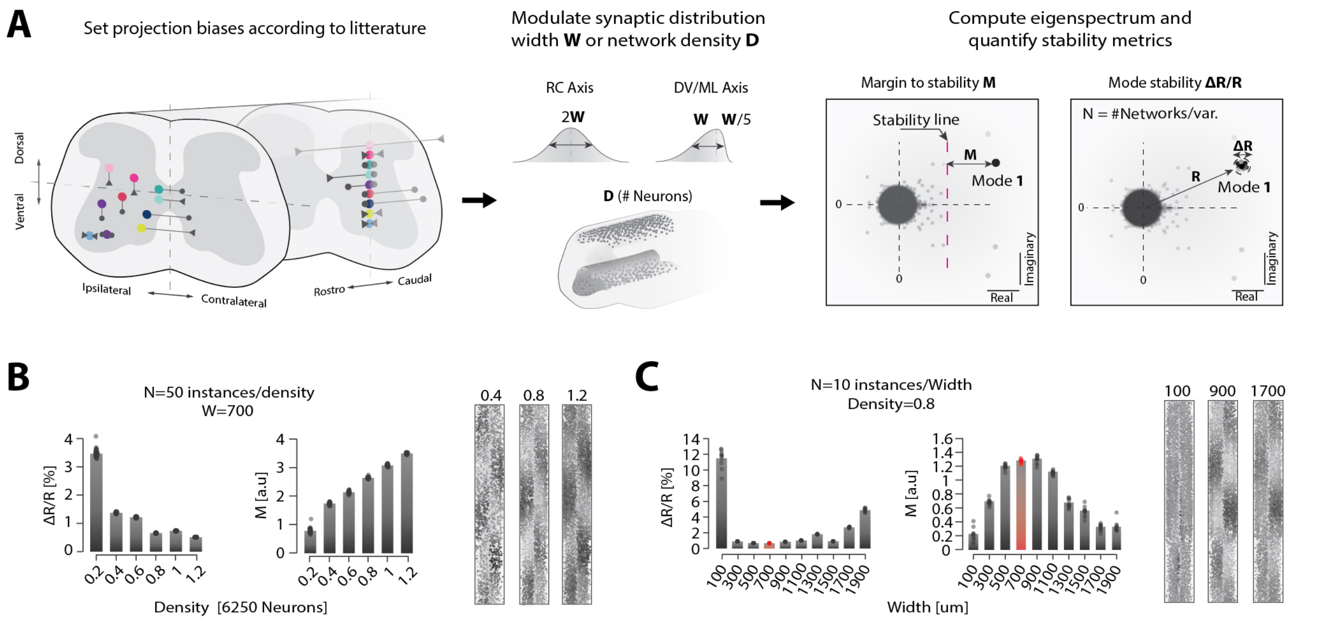


**Fig. S12.** | **Impact of parameters on model stability and emergent dynamics. A**, Procedure to assess model stability. We enforce our choices of projection length and biases as established in Fig. 3 and Fig. S6. We modulate independently two open parameters of the model (synaptic distribution width or network density) and generate several model instantiations for the same set of parameters. For each set of parameters, we compute two metrics on the eigenspectrum of network, reflecting the stability of the dominant mode. (M, distance of the dominant mode to the stability line and ΔR/R, the variability of the dominant mode position. **B**, Quantification of stability metrics for increasing densities (left). Linearly predicted modes for 3 densities (right). While the variability of the dominant mode decreases, its margin to stability increases and the linearly predicted mode is preserved. **C**, Quantifications for increasing projection width (left) and associated modes. There is a optimal width, maximizing the margin to stability and minimizing mode variability (highlighted in red). Instantiations of three projection widths shown right (100, 900, and 1700 $\mu$m, right).
